# “Embryo-eggshell interaction counteracts chiral bias in early *Drosophila* morphogenesis”

**DOI:** 10.64898/2026.03.25.714261

**Authors:** Giulia Serafini, Maryam Setoudeh, Marina B. Cuenca, Charlène Brillard, Matthias Arzt, Pavel Mejstřik, Pierre A. Haas, Pavel Tomančák

**Author notes:** These authors contributed equally (Maryam Setoudeh, Marina B. Cuenca). Pasteur Institute Paris, 25-28 Rue du Dr Roux, 75015 Paris, France (Giulia Serafini). European Molecular Biology Laboratory Barcelona, Carrer del Doctor Aiguader 88, 08003 Barcelona, Spain (Marina B. Cuenca).

## Abstract

Morphogenetic processes during animal development are remarkably invariant (Duboule, 1994; Hall, 1997; Kalinka et al., 2010; Raff, 1996). This stability is established by the interaction between genetic determination of developmental progression and the constraints imposed by the surrounding embryonic environment (Busby and Steventon, 2021; Gilmour et al., 2017; Gorfinkiel and Martinez Arias, 2021). We discovered that the germ band extension process in *Drosophila* is rather variable: instead of extending straight towards the head, the germ band tends to twist to the side. Through a combination of experiments and theory, we demonstrated that Scab integrin-mediated attachment to the vitelline envelope stabilizes the germ band and supports its straight extension. Our quantification of germ band extension dynamics also revealed a consistent handedness to the twist of the germ band. We showed that this left-right asymmetry can be altered by manipulating the expression of Myo1D, the molecular determinant of chirality in *Drosophila* (Lebreton et al., 2018). Our data thus suggest that Myo1D expression causes the early gastrulating blastoderm epithelium to already exhibit inherent chirality and that the resulting destabilization of germ band extension is suppressed by Scab-mediated friction between the blastoderm and the vitelline envelope.

## Introduction

Morphogenetic movements during insect gastrulation follow a stereotyped sequence that is remarkably reproducible across embryos of the same species. Germ band extension in *Drosophila* is an example of this invariant morphogenetic progression (Campos-Ortega and Hartenstein, 1985). A combination of cell intercalations and divisions drives the elongation of the blastoderm tissue in the anteroposterior direction, resulting in the formation of the germ band. The germ band extends around the posterior pole of the embryo (Gheisari et al., 2020; Stern et al., 2022) and on the dorsal side of the embryo it continues to extend towards the head (Campos-Ortega and Hartenstein, 1985). Recently, it has been shown that during this process, the blastoderm interacts mechanically with the vitelline envelope, the innermost layer of the eggshell, in the posterior dorsal region of the embryo (Bailles et al., 2019; Münster et al., 2019). This interaction is mediated by the α integrin subunit Scab, which creates an area of high friction between the cells and the vitelline envelope and is involved in the morphogenesis of posterior midgut invagination (Bailles et al., 2019; Collinet et al., 2024, 2023). In addition to defective posterior midgut invagination, *scab* loss-of-function embryos also show a twisted gastrulation phenotype, in which the germ band twists like a corkscrew around the anteroposterior axis of the embryo (Münster et al., 2019; Smits et al., 2023). This phenotype thus leads to a symmetry breaking along the left-right (L/R) axis of the embryo by the end of stage 8.

By contrast, the first morphological manifestation of L/R asymmetry in wild-type *Drosophila* embryos occurs at stage 13, when the posterior hindgut acquires its characteristic hooked shape (Hayashi and Murakami, 2001; Hozumi et al., 2006a). This chiral deformation has a definite handedness: the hindgut rotates counter-clockwise to position the curve of the gut on the left side in the vast majority of embryos (Hayashi and Murakami, 2001). The key molecule responsible for determining organismal chirality in *Drosophila* is Myosin31DF (Myo1D) (Lebreton et al., 2018; Morgan et al., 1994; Petzoldt et al., 2012a; Spéder et al., 2006). Myo1D exerts forces on the cell cortex by interacting with the actin cytoskeleton and the adherens junctions, establishing planar cell chirality, which is transferred to the tissue level (Chougule et al., 2020; Hozumi et al., 2006b; Lebreton et al., 2018; Petzoldt et al., 2012a; Taniguchi et al., 2011).

Here, we therefore sought to analyse the mechanical and molecular basis for the much earlier L/R symmetry-breaking event represented by the twisting of the *scab* loss-of-function embryos and to understand the role of *scab-*mediated attachment in the overall stability of *Drosophila* germ-band morphogenesis.

Focusing explicitly on L/R symmetry during germ band extension, we discovered that this morphogenetic process is inherently unstable even in wild-type embryos. Through a combination of experiments and mathematical modeling, we demonstrate that Scab-mediated attachment is required throughout germ band extension to stabilize the germ band. Moreover, analysis of the twisting phenotype across different genetic backgrounds revealed a slight but consistent bias towards left-handed twisting. We provide evidence that the determinant of embryonic chirality, Myo1D, can destabilize the germ band during its extension and induce this left-handed twist. Altogether, our data suggest that the early *Drosophila* embryo is chiral long before the obvious morphological manifestations of L/R asymmetry, and that molecular mechanisms such as *scab-*mediated vitelline friction evolved to counteract this inherent chirality.

## Results

### Germ band extension is an unstable process

The hallmark of early *Drosophila* embryogenesis is the extension of the germ band along the embryo midline towards the head during stages 7–10 (Campos-Ortega and Hartenstein, 1985). This process has been considered invariant, with the tip of the extending germ band following a straight line (Fig. 1A). To quantify the straightness of the extension, we characterized the progression of germ band extension by light sheet microscopy in living wild-type *Drosophila* embryos by tracing the position of the dorsal tip of the germ band. This analysis uncovered unexpected phenotypic variability (Fig. 1B; Suppl. Fig. 1): in 43% of embryos from different genetic backgrounds, rather than extending straight towards the head, the germ band curved away from the longitudinal axis of the embryo (Fig. 1C). Our data thus indicate that germ band extension is a potentially unstable developmental process.

**Figure 1:**
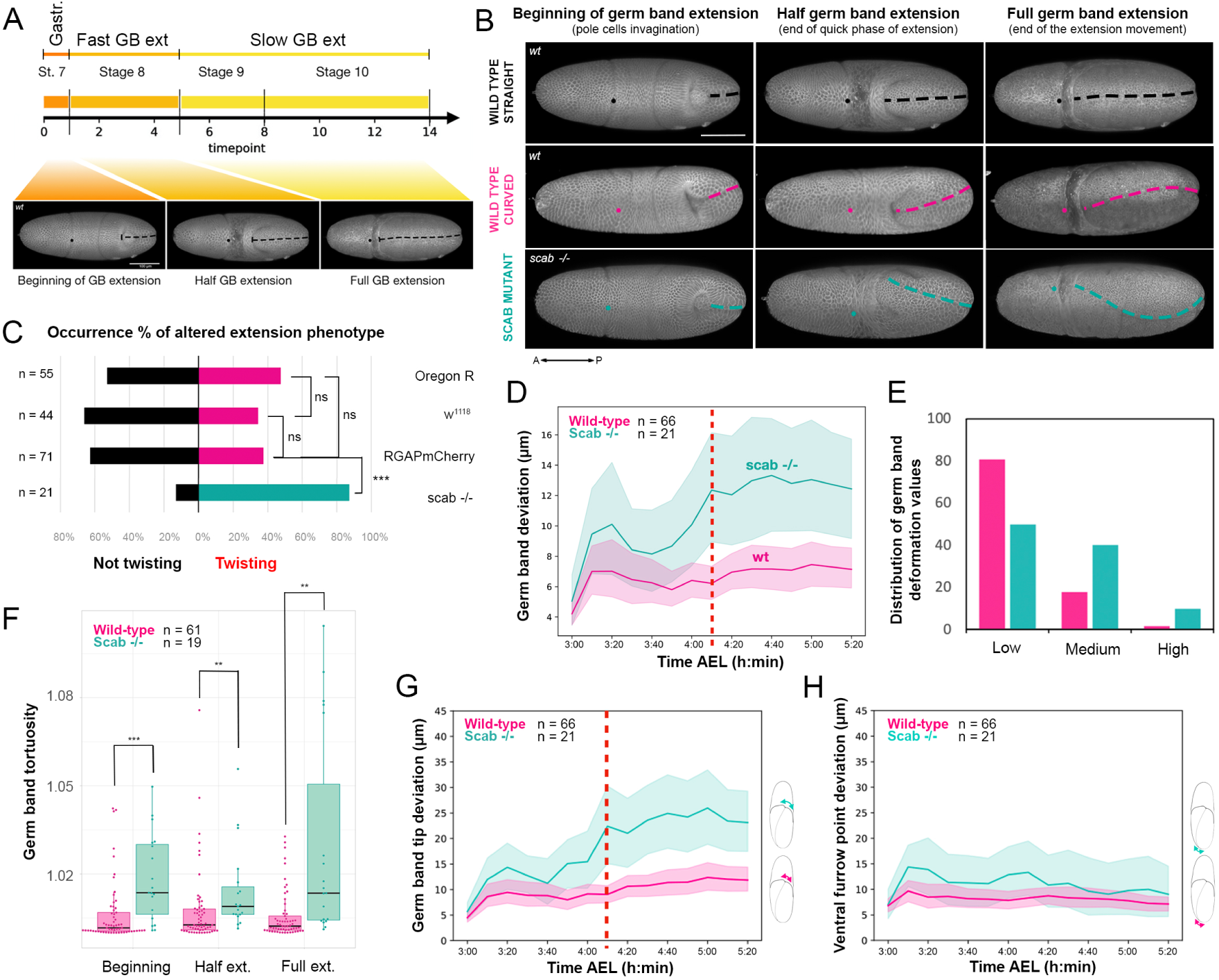
Analysis of germ band extension variability. A) Timeline of the three main phases of germ band (GB) extension and the corresponding developmental stages of *Drosophila*, defining three time points of interest (beginning of GB extension, half GB extension, full GB extension). B) Light sheet recording of wild-type and *scab* mutant embryos at these three time points. Dashed lines highlight the positions of the germ band. Dorsal views. Scale bar: 100 μm. C) Penetrance of the curved germ band phenotype in wild-type genetic backgrounds (magenta *Oregon R, w^1118^, RGAPmCherry*) and in the *scab* mutant (green, *scb^2^/scb^2^* embryos, indicated as *scab^-/-^*). *Oregon R*–*w^1118^*, p > 0.05 (ns); *w^1118^*–*RGAP43::mCherry*, p > 0.05 (ns); *Oregon R*–*RGAP43::mCherry*, p > 0.05 (ns); *RGAP43::mCherry*–*scab^-/-^*, p < 0.001 (***); one-tailed Fisher’s exact test, p-values in Suppl. Table I. D) Plot of the 3D deviation of the germ band midline from a plane (Suppl. Methods) against time in wild-type (magenta, *RGAP43::mCherry/RGAP43::mCherry*) and *scab* loss-of-function (green, *scb^2^/scb^2^*) embryos. The red dashed line indicates the time point after which the difference between the two sample groups becomes statistically significant (p < 0.05). one-way Mann Whitney test, p-values in Suppl. Table III. E) Distribution of germ band deviation values at the end of germ band extension in wild-type (magenta, *RGAP43::mCherry/RGAP43::mCherry*) and *scab* loss of function (green, *scb^2^/scb^2^*) embryos, binned into Low, Medium, High deformation categories for visualization purposes. F) Boxplot of the distribution of germ band tortuosity values at the three time points of interest in the wild-type (*RGAP43::mCherry/RGAP43::mCherry)* and *scab* loss-of-function (*scb^2^/scb^2^*) embryos: p < 0.001 (***); p < 0.01 (**). One-way Mann Whitney test, p-values in Suppl. Table II. G) Plot of the deviation of the tip of the germ band against time in wild-type (magenta, *RGAP43::mCherry/RGAP43::mCherry*) and *scab* loss of function (green, *scb^2^/scb^2^*) embryos. The red dashed line indicates the time point after which the difference between the two sample groups becomes statistically significant (p < 0.05). One-way Mann Whitney test, p-values in Suppl. Table IV. H) Plot of the deviation of the ventral furrow point against time in wild-type (magenta, *RGAP43::mCherry/RGAP43::mCherry*) and *scab* loss of function (green, *scb^2^/scb^2^*) embryos. The two samples are never significantly different. One-way Mann Whitney test, p-values in Suppl. Table V.

We previously observed and reported a similar “twisted gastrulation phenotype” in embryos mutant for Scab integrin, which is proposed to mediate blastoderm-vitelline envelope interaction at the dorso-posterior side of the embryo, required to stabilize gastrulation in *Drosophila* (Bailles et al., 2019; Collinet et al., 2024; Münster et al., 2019). Given the variability in germ band extension observed in wild-type embryos (Fig. 1B, C; Suppl. Fig. S1A, B), we reassessed germ band progression in the *scab*^2^ mutant quantitatively (Fig. 1B-H; Suppl. Fig. S2). We represented germ band deformation using the tortuosity metric, which measures curvature along the midline in 2D (Fig. 1F; Suppl. Fig. S2A–C; Materials and methods for more details). The penetrance of the twisting phenotype was twice as high in *scab* than in wild-type embryos (86% and 39% respectively, Fig. 1C), corroborating the notion that Scab-mediated attachment is required for the germ band to extend in a straight line.

We next asked whether the properties of the twisting phenotype differ between *scab* mutant and wild-type embryos. To that end, we quantified the “deviation from straightness” of the germ band in three dimensions over time by calculating the 3D deviation of the germ band from a 2D plane of lateral symmetry approximating the embryo midline (Fig. 1D; Suppl. Fig. S2H, Suppl. Methods). Because the phenotype is variable in both wild-type and *scab* mutant, we binned the imaged embryos into groups of low, medium, and high twisting to represent the differences in both frequency and intensity of the phenotype between the wild-type and *scab* mutant (Fig. 1E). This analysis shows that the extent of deviation from straightness in the *scab* mutant is greater than in the wild-type, and the frequency shifts towards more extreme twists in the *scab* mutant background. To decompose the complex germ band shape progression, we also separately followed the positions of the two ends of the germ band (Fig. 1G, H): the point where the ventral midline emerges on the dorsal side of the embryo (Fig. 1H, Suppl. Fig. S2F) and the tip of the germ band (Fig. 1G, Suppl. Fig. S2E). The deviation at the point of ventral emergence does not differ significantly between *scab* mutants and the wild-type (Fig. 1H). By contrast, at the germ band tip, the deviation in the *scab* mutant is larger than in the wild-type (Fig. 1G). The change of the deviation from straightness over time (Fig. 1D) corroborates this observation: there is a significant difference between wild-type and *scab* loss-of-function embryos only in the second half of the process, corresponding to the slower phase of germ band extension (Mann Whitney test; p-values in Suppl. Table III; Fig. 1D). Even when focusing on the most altered germ band extension events, germ band deformation is similar between wild-type and *scab* loss of function embryos in the first timepoints, while it diverges later on (Fig. 1D, Suppl. Fig. S2D). In summary, our spatiotemporal analysis of the germ band deviations suggests that common sources of instability act on the germ band from the earliest stages in both wild-type and *scab* mutants, and that this instability is counteracted by Scab at later stages.

To further support the role of Scab integrin in the stabilization process, we imaged the expression and subcellular localization of the Scab integrin subunit throughout germ band extension using a transgenic line expressing Scab::mNeonGreen (Scab::mNG) under the control of the *scab* endogenous promoter. Light sheet imaging revealed Scab::mNG protein in the posterior dorsal region of the cellular blastoderm as early as stage 5 of embryogenesis (Fig. 2A; Suppl. Fig. S3). The protein was then continuously expressed in cells anterior to the tip of the extending germ band until the end of the process (Fig. 2A,B; Suppl. Fig. S3). Quantification of spatial and temporal protein expression dynamics showed that Scab expression begins in a broader domain at relatively low intensity. The expression area and intensity initially decrease, presumably due to reorientation of blastoderm cells during hindgut invagination. Later, this domain expands and signal intensity increases steadily, remaining high throughout germ band extension (Fig. 2D). At the subcellular level, early on during the fast germ band extension phase, Scab protein is predominantly localized apically in epidermal cells. Later, this apical localization persists, but the protein is also detected throughout the epidermal cells (Fig. 2B,C,E). Throughout germ band extension, apical localization is characterized by distinct foci of Scab signal at the cell surface (Fig. 2B,C). The foci were particularly prominent at the point of last contact between cells and the vitelline envelope, before detachment due to bulk tissue movement and the analysis of time-lapse images revealed that the Scab foci persist after detachment (Fig. 2B). This correlation of foci with cell-vitelline envelope detachment was observed throughout germ band extension. To connect Scab expression with cell attachment to the vitelline envelope, we performed transmission electron microscopy analysis of the apical region of cells expressing Scab (Fig. 2F). At both early and late stages of germ band extension, we detected contacts between Scab-positive cells and the vitelline envelope and/or the extracellular matrix material of the perivitelline space. EM analysis revealed that areas where cells contact the vitelline envelope exhibit greater ECM deposition in the perivitelline space. Moreover, the cells in contact with the ECM/vitelline have rough contours, supporting the idea of friction between the vitelline envelope and cells expressing *scab*.

**Fig. 2:**
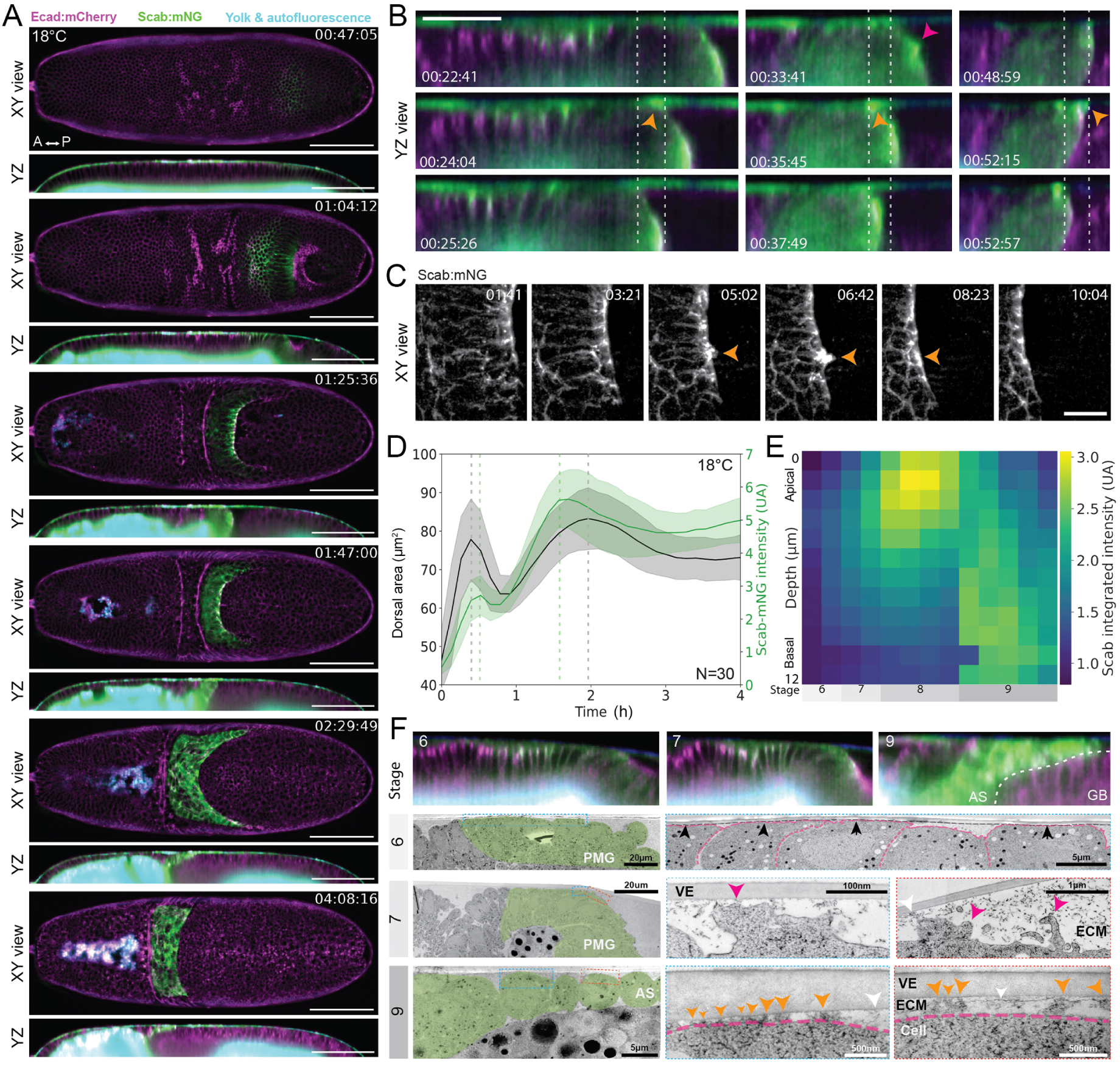
Scab protein expression. A) Time lapse of germ band extension in an embryo expressing the E-Cadherin:mCherry (magenta) and Scab:mNG (green) live reporters imaged at 18 °C. Dorsal views and local Z-projections along the X axis are shown for each of the selected timepoints. The autofluorescence of the yolk and vitelline membrane is shown in cyan. Scale bar: 100 μm. B) Close-up views of the dynamics of attachment and Scab-foci (arrowheads) at three different time points (columns). YZ view (intensity scale adjusted to prevent saturation). Scale bar: 50 μm. C) Timelapse of Scab protein expression pattern during hindgut invagination, showing repartitioning at the very edge of the invagination. D) Plot of the dorsal area (gray curve) and the intensity (green curve) of the domain of expression of the Scab:mNG protein against time (N = 30). E) Heat map of the integrated Scab intensity at different depths against time (developmental stage), illustrating basal relocalization of Scab after a peak in apical recruitment. F) Light (top row) and electron microscopy (EM) snapshots of different tissues and at different stages. Green overlay marks Scab expressing cells in EM based on correlation with live-imaging data from similar stages. Middle and right columns: close-up views of boxed regions in the left column. VE – vitelline envelope, ECM – extracellular matrix. Black arrowheads at stage 6 row show the zones of contact with the VE whose bottom boundary is marked with a black line. Orange and magenta arrowheads at stage 7 and 9 rows point to attached and detached cell protrusions, respectively. White arrowheads at stage 7 and 9 rows point to an electron-dense structure that could represent transmembrane proteins and filamentous ECM structures under the VE.

In summary, the observed Scab expression is consistent with the hypothesis that Scab-mediated attachment counteracts an instability of germ band extension as the germ band progresses toward the anterior of the embryo. We next turned to biophysical modeling to understand the sources of this instability.

### A mechanical model of germ band extension predicts germ band shape variability

To study the forces at play during germ band extension and understand the mechanical basis for germ band shape variability, we derived a model that reduces the mechanics of the dorsal side of the germ band, which is a bulk tissue, to effective mechanics of the germ band midline (Fig. 3A). In our simplified model, the germ band midline is an elastic line that is pushed by a force *F* representing germ band extension (Fig. 3B). The elasticity of the midline resists its bending, while the surrounding tissues and the vitelline envelope generate friction forces that resist motion perpendicular and parallel to the direction of germ band extension, with associated friction coefficients *γ*_⊥_, *γ*_∥_ (Fig. 3B). Scab-mediated attachment at the tip of the germ band generates additional friction forces there, with corresponding coefficients *ζ*_⊥_, *ζ*_∥_ in the perpendicular and parallel directions, respectively (Fig. 3B).

**Figure 3:**
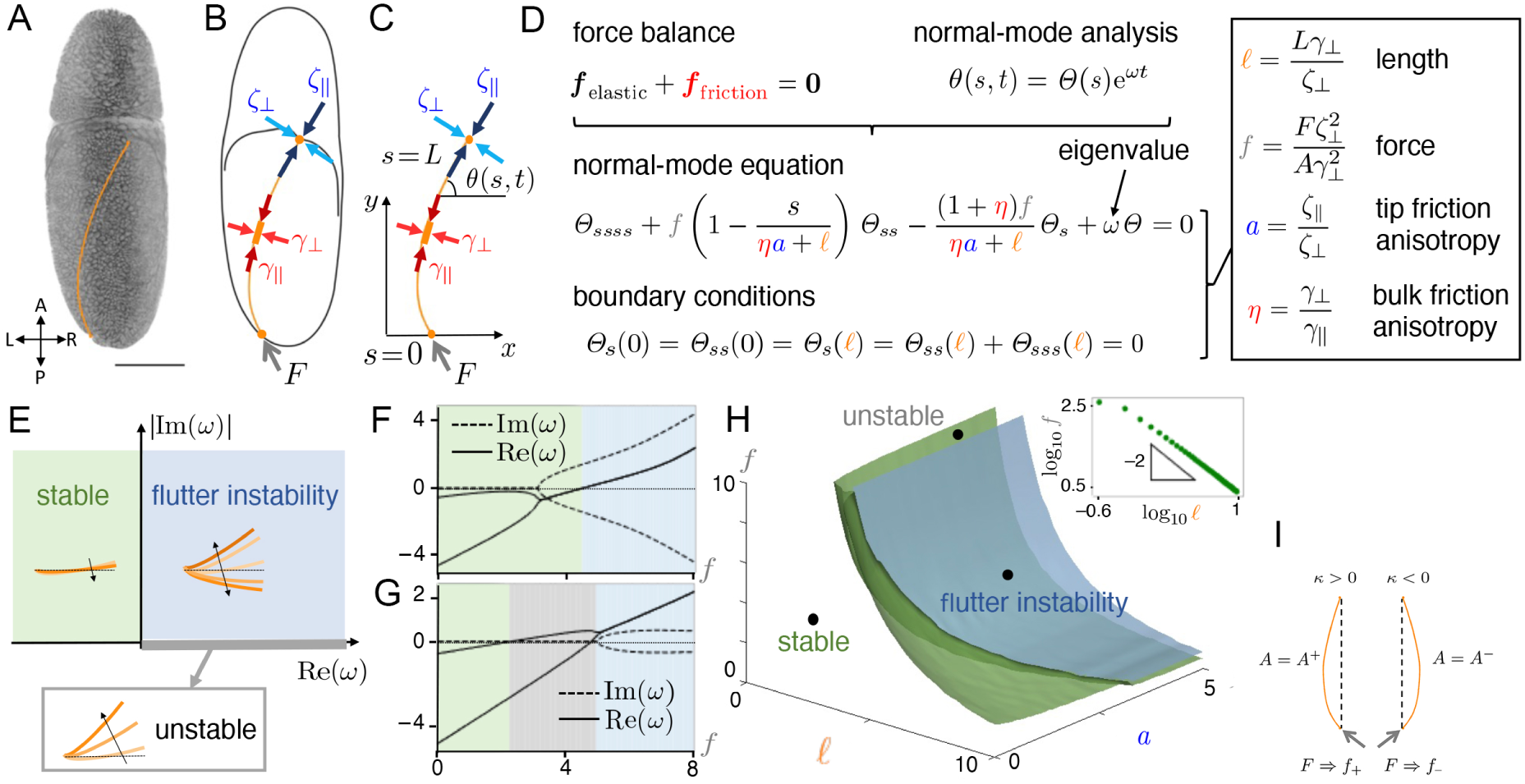
Mathematical model of germ band extension. A) Dorsal view of a *Drosophila* embryo during germ band extension. Orange line: germ band midline. Embryo axes: A(nterior), P(osterior), L(eft), R(ight). Scale bar: 100 μm. B) Mechanical model of the germ band midline as an inextensible elastic line pushed by a force *F* driving the extension. Bulk forces along the germ band midline with perpendicular and parallel friction coefficients *γ*_⊥_, *γ*_∥_ represent friction forces from the surrounding tissues and the vitelline envelope. Tip forces with perpendicular and parallel friction coefficients *ζ*_⊥_, *ζ*_∥_ represent friction from the tip attachment of the germ band. C) Two-dimensional projection of the model into the (*x*, *y*) plane. Arclength along the inextensible germ band midline is *s*, with *s* = *0* corresponding to the posterior end and *s* = *L* to the tip of the germ band. The tangent angle to the germ band is *θ*(*s*, *t*). D) Governing equations of the mechanical model: balancing the elastic and friction forces in the bulk and making the normal mode ansatz *θ*(*s*, *t*) = *Θ*(*s*) e*^ωt^* leads to a fourth-order differential problem for the (complex) eigenvalue *ω*. This problem depends on four dimensionless parameters: the dimensionless germ band length *ℓ*, the dimensionless pushing force *f* depending on the bending modulus *A* of the germ band midline, the tip friction anisotropy *a*, and the bulk friction anisotropy *η*. E) Interpretation of the eigenvalue *ω* in terms of its real part Re(*ω*) and the absolute value of its imaginary part Im(*ω*): if Re(*ω*) < 0, small perturbations to the straight shape decay and the germ band is stable (green shaded area); if Re(*ω*) > 0 and Im(*ω*) = 0, such perturbations grow monotonically, corresponding to an unstable germ band (grey shaded line); if Re(*ω*) > 0 and Im(*ω*) ≠ 0, such perturbations grow in an oscillatory manner that corresponds to a “flutter” instability (blue shaded area). F) Plot of Re(*ω*) and Im(*ω*) against *f* for the two leading eigenvalues for *ℓ* = 5, *a* = 1.01, *η* = 2. For small *f*, the germ band is stable; for larger *f*, there is a flutter instability. G) Corresponding plot for *a* = 12.9. The germ band is unstable in an intermediate range of *f*. H) Phase diagram of the instability in (*ℓ*, *a*, *f*) space for *η* = 2, indicating the regions of parameter space in which the germ band is stable, unstable, and shows a flutter instability, respectively. Inset: plot of the critical value of *f* against *ℓ* for fixed *a* = 0.2 on logarithmic axes, showing the approximate scaling *f* ∝ *ℓ*^−2^. I) Asymmetry of the bending modulus *A* depending on the curvature *κ*, viz., *A* = *A*_+_ if *κ* > 0, *A* = *A_–_* if *κ* < 0, leads to asymmetry of the dimensionless pushing forces, *f_+_* for *κ* > 0, *f_–_* for *κ* < 0, resulting from the same physical pushing force *F*.

We simplified the curved geometry of the tissue by projecting the dorsal view of the germ band midline into the plane, describing the shape of the midline by its tangent angle *θ*(*s*, *t*), in which *s* ∈ [0, *L*(*t*)] is arclength along the germ band midline, with *L*(*t*) the length of that midline that increases with the time *t* of germ band extension. We assume the midline to be elastically inextensible (Fig. 3C). We impose the balance between friction forces and elastic forces resulting from bending of the midline and tension in the midline. We describe the shape of the germ band midline in this model using normal modes *θ*(*s*, *t*) = *Θ*(*s*) e*^ωt^*, and we obtain a differential equation for the shape *Θ*(*s*) and the eigenvalue *ω* in the Supplemental Note and Fig. 3D. This differential equation depends on four dimensionless parameters defined in Fig. 3D: the dimensionless germ band length *ℓ*, the dimensionless pushing force *f*, the anisotropy *a* of the tip friction coefficients, and the anisotropy *η* of the bulk friction coefficients. The eigenvalue *ω* reveals the stability of the germ band (Fig. 3E): if Re(*ω*) < 0, then small perturbations to a straight germ band midline decay and the midline remains straight (Fig. 3E, green area of the graph). If Re(*ω*) > 0 and Im(*ω*) = 0, then such perturbations grow monotonically. This unstable case corresponds to the observed germ band variability (Fig. 3E, grey part of the x-axis). If, however, Re(*ω*) > 0 and Im(*ω*) ≠ 0, then there is “flutter instability”: small perturbations to a straight midline grow in an oscillatory manner that is not observed in experiments (Fig. 3E, blue area of the graph). Determining *ω* numerically (Supplemental Note), we found that the germ band midline is stable if the pushing force parameter *f* is small enough and that there is flutter instability for large *f* (Fig. 3F), while it is possible for the germ band to be unstable in an intermediate range of *f* depending on the values of the parameters *ℓ*, *a* (Fig. 3G). This showed that the model can explain the observed curved extension of the germ band midline. More quantitatively, we summarise the results of our calculations in a bifurcation diagram (Fig. 3H) that shows the regions of parameter space (*ℓ*, *a*, *f*) in which the different stability outcomes arise; this does not change qualitatively with the bulk friction anisotropy parameter *η* (Supplemental Note). At constant germ band length *ℓ*, the minimum force required for instability decreases with increasing tip friction anisotropy coefficient *a* (Fig. 3H). Similarly, at constant *a*, this critical force decreases with increasing *ℓ* with the approximate scaling *f*_crit_ ∝ *ℓ*^−2^ (Fig. 3H, inset) expected from classical Euler buckling (Landau and Lifshitz, 1986). These results allowed us to understand the twisting phenotype displayed by the *scab* loss of function mutants. Indeed, we expect removal of the Scab-mediated attachment to reduce strongly the tip frictions ζ_⊥_, ζ_∥_, and hence, assuming that the *scab* mutation does not affect the other model parameters, to increase *ℓ* and decrease *f* in such a way that *f ℓ*^2^ remains constant (Fig. 3D). Because of the observed scaling of the stability boundary *f*_crit_ *ℓ*^2^ = const. (Fig. 3H, inset), this means that the model predicts that the increased twisting of the *scab* mutants requires *a* to be larger in the mutant than in the wildtype. In turn, this implies that *scab* depletion reduces the twisting friction ζ_⊥_ more strongly than the friction ζ_∥_ resisting germ band extension. This is consistent with the idea that germ band twisting is specifically resisted by Scab attachment, while the extension movement is also opposed by the interaction with the extraembryonic tissue of the amnioserosa.

The cell intercalations driving germ band extension lengthen the germ band midline and generate effective forces at both ends of the midline (Stern et al., 2022). By representing germ band extension as a pushing force, our model assumes that the mechanics of germ band extension are dominated by these forces rather than its lengthening. Representing the germ band as a pushed line is also in agreement with recent results from Stern et al. (2022), who tracked the localization of the T1 and rosette transitions on a single cell level and showed that the rosette transitions involve mostly cells located ventrally to the equator of the embryo and towards the posterior region, while the T1 transitions involve cells uniformly distributed on the lateral sides of the embryo (Stern et al., 2022). When considering the embryo from the dorsal side, the sum of these movements can be approximated to a force pushing the germ band from the posterior of the embryo, which is the pushing force applied on our elastic line. Another way of representing germ band extension dynamics would be to account more explicitly for the effect of the cell divisions occurring in the various mitotic domains along the germ band rather than the effect of cell intercalations (Foe, 1989). To test this hypothesis, we analyzed the opposite model, in which the germ band midline is a slowly growing elastic line, but there is no pushing force. We found (Supplemental Note) that this model allows only stability and flutter instability, hence cannot explain the observed curved extension. This supports our hypothesis and suggests that the “pushed” model recapitulates better the mechanics *in vivo*. Neither of these models includes the curvature of embryonic surface, however; we briefly discuss the mathematical description of this effect in the Supplemental Note.

### Left-right asymmetry in the early stages of embryogenesis

The germ band twisting phenotype inherently breaks the left-right symmetry of the embryo. We thus asked whether the twist to the left or right is random. To increase the throughput of this analysis, we examined fixed embryos. We used the *ind* (*intermediate neuroblast defective*) expression pattern as an unambiguous marker of the handedness of the germ band twist in the analysis of fixed samples (Fig. 4A). The results revealed an unexpected bias in twisting directionality, with an average of 67% of embryos having the germ band tip directed to the left (Fig. 4C). We replicated this result in live imaging experiments, in which the dynamics of germ band extension were observed (Fig. 4B, D). This bias was always towards the left side, was independent of the balancer’s genetic background, manifested in fixed as well as live imaged embryos, and extended even to a panel of deficiencies previously shown to exhibit curved germ band extension (Bonds et al., 2007) (Fig. 4E, F, and Suppl. Fig. S4).

**Figure 4:**
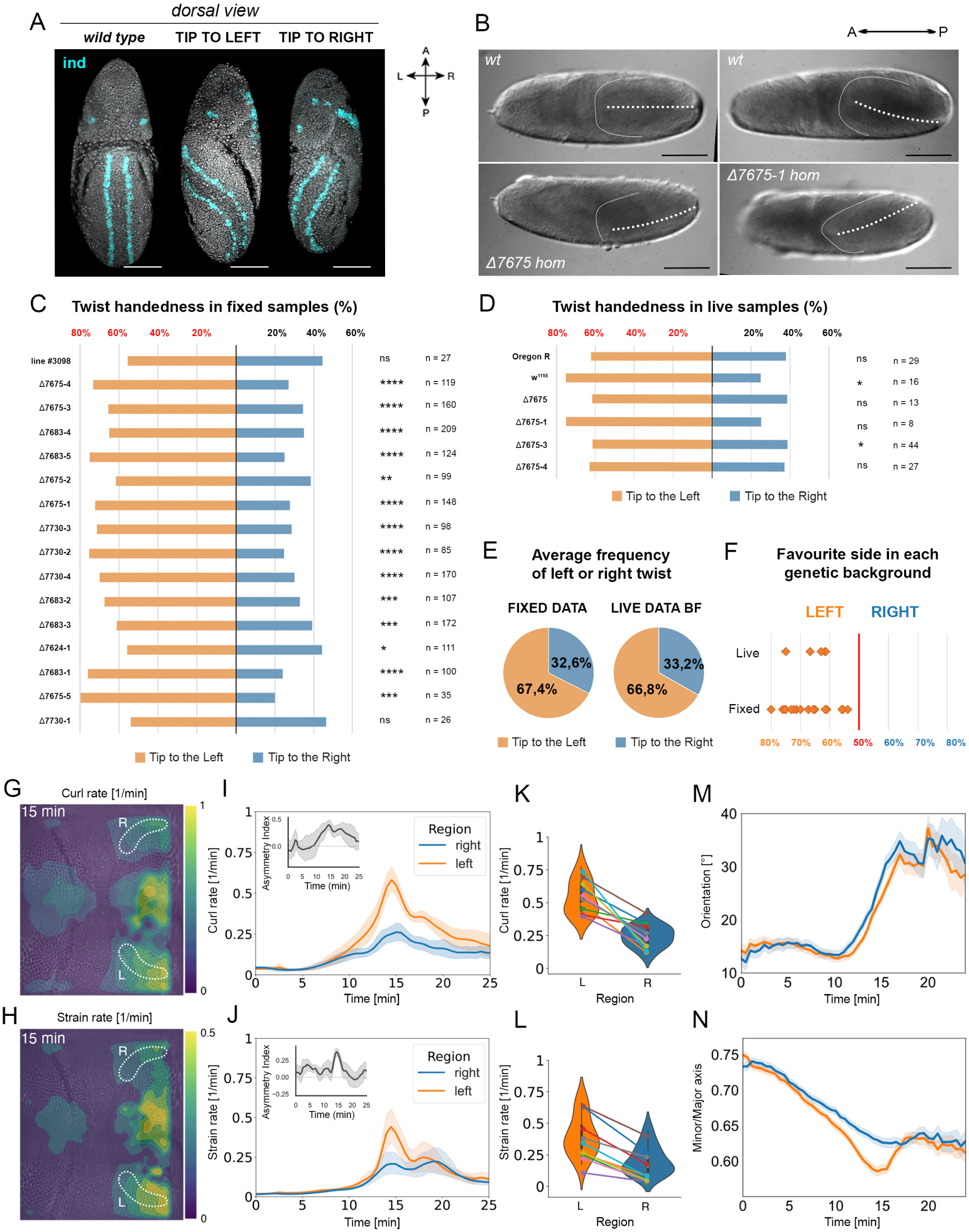
Left-right asymmetry in the early Drosophila embryo. A) Example of an *ind* pattern (cyan) in embryos showing straight, twisted to the left, and twisted to the right germ band extension. Dorsal views. Scale bar: 100 μm. B) Brightfield recording of embryos of different genetic backgrounds showing altered germ band extension. Genotypes in Suppl. Table XII. Dorsal views. Scale bar: 100 μm. C) Distributions of the handedness of the twists in fixed samples of various genotypes (scored as illustrated in panel A). Orange: twist towards the left side; blue: twist towards the right side. p < 0.0001 (****); p < 0.001 (***); p < 0.01 (**); p < 0.05 (*); p > 0.05 (ns). Binomial test, p-values in Suppl. Table VI and genotypes in Suppl. Table XII. D) Distributions of the handedness of the twists in live samples derived from the brightfield imaging experiments (as illustrated in panel B). Orange: twist towards the left side; blue: twist towards the right side. p < 0.0001 (****); p < 0.001 (***); p < 0.01 (**); p < 0.05 (*); p > 0.05 (ns). Binomial test, p-values in Suppl. Table VI and genotypes in Suppl. Table XII. E) Pie-chart representing the average frequency of twist handedness in the fixed and the live brightfield data (panels C, D). Orange: twist to the left; blue: twist to the right. F) Preferred handedness of germ band twisting in each of the lines analyzed (panels C, D). Each dot represents the bias of twist in a different genotype. G) Magnitude of curl rate at *t* = 15 minutes after cellularization, when the asymmetry between the left and right sides is largest (corresponding to the maximum in panel I). The ROIs analyzed are bounded by the white dotted lines. H) Magnitude of strain rate at *t* = 15 minutes after cellularization, when the asymmetry between the left and right sides is largest (corresponding to the maximum in panel J). The ROIs analyzed are bounded by white dotted lines. I) Plot of the mean values of curl rate on the left (orange) or right (blue) sides of the posterior midgut invagination against time *t*, where *t* = 0 marks the end of cellularization. The inset shows the corresponding asymmetry index for each timepoint. J) Plot of the mean values of strain rate on the left (orange) or right (blue) sides of the posterior midgut invagination against time *t*, where *t* = 0 marks the end of cellularization. The inset shows the corresponding asymmetry index for each timepoint. K) Mean value of curl rate at *t* = 15 minutes, showing a significant difference across embryos (n = 10; p < 0.001, one-sided Wilcoxon signed-rank test, p-values in Suppl Table VII). L) Mean value of strain rate at *t* = 15 minutes, showing a significant difference across embryos (n = 10; p < 0.001, one-sided Wilcoxon signed-rank, p-values in Suppl. Table VII). M) Mean orientation of ellipses fitted to cell shapes against time, for left and right ROIs. No significant difference is observed between the two sides (Wilcoxon signed-rank test). N) Ratio of minor to major axis lengths of the fitted ellipses against time. Lower values, indicating higher deformation, are reached for the cells in the left ROI (p < 0.05, one-sided Wilcoxon signed-rank test, p-values in Suppl. Table VII).

To confirm the bias at the tissue scale, we analyzed tissue flows on the two sides of the germ band in gastrulating embryos imaged with light sheet *in-toto* (Fig. 4G–L; Suppl. Fig. S4). By particle image velocimetry (PIV) analysis, we decomposed the tissue flows into the pure deformation (strain) and pure rotation (curl) components (Fig. 4G–J and Suppl. Fig. S4A, B). We found that both the curl and strain rates are significantly higher on the left side of the embryo (p < 10^−3^; n = 10; one-sided Wilcoxon signed-rank test), reaching their highest asymmetry 15 minutes after the onset of gastrulation (Fig. 4I–L). At the cellular level, we segmented single cells on both sides of the posterior midgut invagination and fitted ellipses to them (Suppl. Fig. S4C). While the dynamics of the orientation of these fitted ellipses are similar on the left and right sides, cells on the left side show a higher deformation, as quantified by the ratio of their minor and major axes (Fig. 4M, N). The difference was again the largest 15 minutes after the start of gastrulation (p < 0.05; n = 10; one-sided Wilcoxon signed-rank test), reflecting the asymmetry quantified at the tissue-level (Fig. 4I, J, and M, N). In summary, the asymmetry in cellular and tissue dynamics is consistent with the observed bias in the handedness of germband twisting (Fig. 4C–F). To investigate the possible origins of this left-right bias of the instability of the germband, we returned to our biophysical model.

### The mechanical model predicts that left-right asymmetry can cause curved extension bias

To link left-right asymmetry to the observed bias of curved extension we postulate that the observed asymmetry of tissue deformation rates and cell shapes (Fig. 4) leads to an asymmetry of the effective bending modulus *A* of the midline depending on the bending direction, i.e., on the curvature *κ* of the midline, viz., *A* = *A*_+_ if *κ* > 0, *A* = *A_–_* if *κ* < 0, and *A*_+_ > *A_–_* without loss of generality (Fig. 3I). At the same physical pushing force *F*, this implies that the dimensionless pushing force *f* takes values *f_+_* for *κ* > 0, *f_–_* for *κ* < 0, where *f_+_* < *f_–_* (Fig. 3D and 3I). It is possible that the midline is unstable for *f_–_*, but is still stable for *f_+_* (Fig. 3G), so that bending towards *κ* < 0 is favored and curved extension is biased. We do not expect the bias resulting from this mechanism to be perfect however, not only due to intrinsic variability: for example, if *f_+_* < *f_–_* are both unstable, with eigenvalues *ω_+_, ω_–_* > 0, it is possible (Fig. 3G) that *ω_+_* > *ω_–_*, i.e., that the instability of the mode with *κ* > 0 grows faster, and so bending in this direction is favored instead.

This argument extends to suggesting that left-right asymmetry favors the instability: if we additionally assume a fixed average bending modulus *Â* of the tissue, then *A* = *Â* in the absence of left-right asymmetry, and *A*_+_ > *Â* > *A_–_*. A similar argument shows that the dimensionless pushing force *f̂* in the symmetric case satisfies *f_+_* < *f̂* < *f_–_*, so the midline can be unstable in the left-right asymmetric case while being stable in the absence of left-right asymmetry.

### The early presence of molecular chirality could underlie germ band extension instability

A possible mechanistic explanation for the observed left-right (L/R) bias in germ band twist lies in the molecular determinants of cell chirality. The earliest developmental event in *Drosophila* that exhibits L/R asymmetry is gut morphogenesis, which begins 9h 20’ after egg laying at stage 13 (Inaki et al., 2016). The key molecule responsible for gut rotation is the unconventional Myosin 1D (Myo1D, also known as Myo31DF); (Petzoldt et al., 2012b). Myo1D was proposed as the determinant of embryonic chirality, as manipulating its expression confers chiral properties to the tissues in which it is expressed (Lebreton et al., 2018).

During early embryogenesis, the *myo1D* transcript is deposited maternally, and its zygotic expression pattern is dominated by strong upregulation in the presumptive hindgut region at stage 6 (Suppl. Fig. 5A, B); (Tomancak et al., 2007, 2002). During germ band extension, *myo1D*-expressing cells are internalized and form the hindgut primordium. We visualized Myo1D protein expression during early embryogenesis using live imaging of the Myo1D::mNG reporter (Chougule et al., 2020); (Fig. 5A). At the blastoderm stage, Myo1D protein was localized ubiquitously at the cell membrane. During cellularization, the Myo1D signal followed invaginating membranes basally (Suppl. Fig. 5B, C). After cellularization, throughout germ band extension, Myo1D signal was ubiquitously detected at the boundaries of epidermal cells (Fig. 5B). At organogenesis stages, the Myo1D signal peaks in the hindgut (Fig. 5C). Quantification of the Myo1D protein signal (Fig. 5D) shows that the maternal pool of protein is strongly reduced at the time of posterior midgut invagination, suggesting that zygotic expression of Myo1D occurs in the germ band. Thus, the chiral determinant Myo1D is present in germ band cells during L/R-biased curved extension.

**Figure 5:**
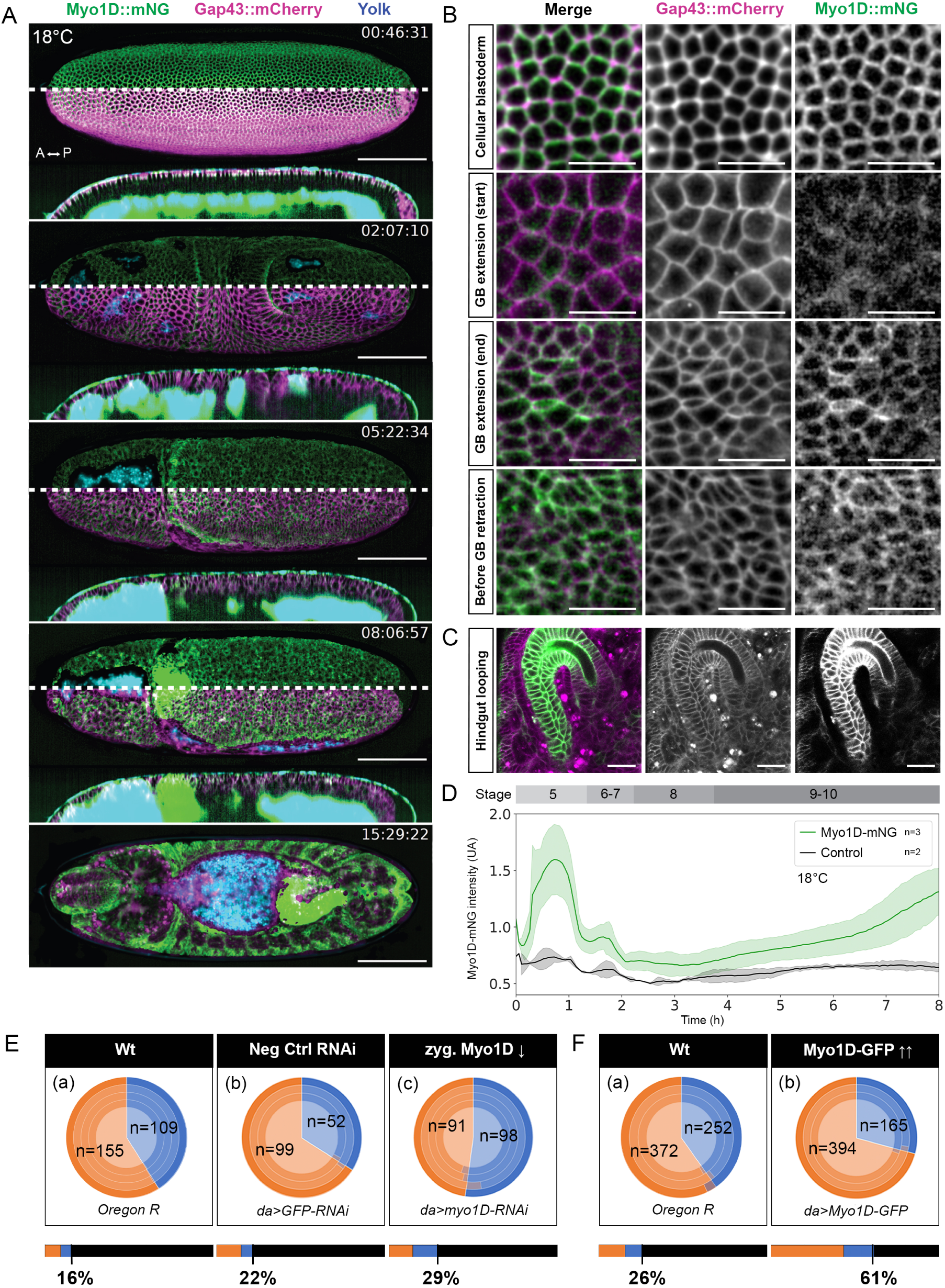
Myo1D function in early *Drosophila* embryogenesis. A) Live expression pattern of Myo1D::mNG (green) in the presence of the fluorescent membrane marker RGAP43-mCherry (magenta) at selected timepoints representative of different phases of germ band extension. The autofluorescence of the yolk and vitelline envelope is indicated in blue. Dorsal views and longitudinal cross-sections. Scale bar: 100 μm. B) Live expression of Myo1D::mNG reporter in the presence of RGAP43::mCherry membrane marker at different stages: in the cellular blastoderm at the cell boundaries and throughout the extension of the germ band. Scale bars: 20 μm. C) Live expression of Myo1D::mNG reporter in the presence of RGAP43::mCherry membrane marker in the hindgut. Scale bar: 20 μm. D) Expression levels of the Myo1D::mNG reporter throughout the first 8 hours of embryonic development (green line) compared to the background signal in a wild-type embryo (black line). Scale bar: 100 μm. E) Effects on germ band extension handedness bias and germ band twist penetrance in conditions of *myo1D* knock-down. The pie charts indicate the germ band handedness of the embryos performing twisted germ band extension (orange: twist left and blue: twist right). Each lighter circle is an individual replicate; the darker circle shows the average value: (a) wild-type embryos, (b) negative control embryos, (c) embryos in which *myo1D* expression was downregulated. The detected handedness bias is statistically significant for each replicate (Binomial test; p-values in Suppl Table VIII), and the leftward bias is significantly reduced upon *myo1D* knockdown (Binomial Generalized Linear Mixed Models (GLMMs); p-values and Odds Ratio Percentages in Suppl. Table IX). The bars under the pie charts indicate the twist penetrance for each genotype (orange: twist left, blue: twist right, black: no twist): (a) wild-type embryos (n = 1645), (b) negative control of the RNA interference (n = 678), (c) embryos downregulating *myo1D* (n = 644). The germ band twist penetrance in the different samples has been compared using Binomial Generalized Linear Mixed Models (GLMMs) (p-values and Odds Ratio Percentages in Suppl. Table X). F) Effects on germ band extension handedness bias and germ band twist penetrance in conditions of Myo1D::GFP overexpression. The pie charts are labeled as in (E). (a) wild-type embryos, (b) embryos overexpressing Myo1D::GFP maternally. The detected handedness bias is statistically significant for each replicate (Binomial test; p-values in Suppl Table VIII) and the handedness of the samples has been compared using Binomial Generalized Linear Mixed Models (GLMMs) (p-values and Odds Ratio Percentages in Suppl. Table IX). The bars under the pie charts indicate the twist penetrance for each genotype as in (E). (a) wild-type embryos (n = 2367), (b) embryos overexpressing Myo1D::GFP protein (n = 911). There is an average 4-fold increase in twist penetrance in embryos overexpressing Myo1D::GFP (Binomial Generalized Linear Mixed Models (GLMMs); p-values and Odds Ratio Percentages in Suppl. Table X).

Since germ band twisting is strongly enhanced in *scab* mutants and *scab* is expressed in a similar area of the blastoderm as *myo1D*, we assessed the spatiotemporal relationship between the Scab and Myo1D proteins through live imaging of reporter lines (Suppl. Fig. S5B). The results showed that, although the expression patterns of the two genes partially overlap at the blastoderm stage, after germ band extension, the cells expressing *scab* and *myo1D* are distinct. Scab protein was expressed in the amnioserosa, while the Myo1D protein was expressed at the tip of the germ band, in the cells of the hindgut primordium (Fig. 5A–C). On the other hand, we observed that the Myo1D protein is ubiquitously expressed in the germ band epidermis during germ band extension. Its signal increases after posterior midgut invagination, and throughout the slow phase of germ band extension when *scab* mutants exhibit the pronounced twist (Fig. 1B, D; Fig. 5A, D). We therefore hypothesized that the amount of Myo1D present in the germ band epidermis could influence its extension by imparting cell chirality.

### Altering Myo1D functionality affects embryonic chirality

To test whether the Myo1D protein could destabilize germ band extension and bias the directionality of its twist, we studied germ band extension in conditions of altered Myo1D function (Fig. 5E, F).

We knocked down *myo1D* expression using *Gal4-*driven RNA interference to impede zygotic Myo1D production (Fig. 5Ea–c). Upon RNA interference, *myo1D* zygotic expression was depleted at least 3.5-fold compared to wild-type intensity (∼71% signal reduction, two-sample T-test, p-values in Suppl. Table XI; Suppl. Fig. S6A–C). In agreement with our hypothesis, we observed the disappearance of the leftward bias in the samples where Myo1D protein function had been depleted (Fig. 5Ec, *da>myo1D-RNAi*). The negative RNAi control (*da>GFP-RNAi*) showed the same leftward bias as the wild-type (Fig. 5Eb; Binomial test, p-values in Suppl. Table VIII; GLMM analysis, p-values and Odds Ratio Percentages in Suppl. Table IX).

Conversely, we utilized a ubiquitous driver to achieve the maternal overexpression of a Myo1D::GFP construct known to rescue the loss-of-function phenotype (Lebreton et al., 2018) (Fig. 5Fa, b; Suppl. Fig. S6D). Upon Myo1D::GFP overexpression, we observed a four fold increase in the penetrance of the twisted germ band extension phenotype compared to that observed in the wild-type sample. This increase was much bigger than the increase observed in the Myo1D-RNAi and GFP-RNAi samples (Fig. 5F; GLMM analysis, p-values and Odds Ratio Percentages in Suppl. Table X). On the other hand, the handedness of the bias did not change dramatically upon Myo1D::GFP overexpression. The majority of the embryos were still twisting towards the left side and the leftward twisting frequency increased from ∼60% observed in the wild-type sample, to ∼70% in the Myo1D::GFP protein overexpression (Fig. 5F; Binomial test, p-values in Suppl. Table VIII; GLMM analysis, p-values and Odds Ratio Percentages in Suppl. Table IX).

Taken together, the loss-of-function and gain-of-function analyses indicate that the molecular chiral determinant Myo1D confers a leftward bias on germ band extension when the germ band veers off course. Since Myo1D overexpression also enhances the frequency of curved germ band twists, the Myo1D activity may be the underlying cause of the instability of germ band extension.

## Discussion

Our results indicate that germ band extension is inherently mechanically unstable even under normal developmental conditions in wild-type embryos. This agrees with previous observations (Smits et al., 2023). With hindsight, it is perhaps not entirely surprising that the genome-encoded developmental progression cannot fully control such a complex bulk tissue movement in confined spaces. This is confirmed by our minimal mechanical model, in which the opposing forces driving and resisting germ band extension result in a mechanical instability leading to curved extension. A regulatory feedback loop might counter this instability, but it is inherently difficult for the developmental system to detect an instability at the organismal level. Instead, our analysis of the *scab* loss-of-function phenotype suggests that a redundant mechanism stabilizing the germ band progression exists in fruit flies: both the physical model and the expression analysis support the hypothesis that *scab-*mediated attachment of the blastoderm cells to the inner surface of the vitelline envelope counteracts the germ-band instability throughout the extension process. This stabilization mechanism is analogous to the recently proposed role of the cephalic furrow in absorbing compressive stresses at the head-trunk boundary during germ-band extension in *Drosophila* embryos (Dey et al., 2025; Vellutini et al., 2025). In that context, the genome-encoded developmental progression cannot control or detect the tissue collisions arising from the temporal coincidence of two independent morphogenetic processes (mitotic domain divisions and germ band extension) either. There, the solution appears to be the cephalic furrow as a mechanism to relieve the compressive stresses. In both processes, germ-band extension and cephalic furrow formation, these stabilizing mechanisms are not fully effective: the germ band twists slightly, and ectopic folds appear near the cephalic furrow, even in wild-type embryos. Interestingly, the selective abolition of cephalic furrow morphogenesis leads to a twisting germ band, suggesting that multiple independent mechanisms may stabilize the mechanically complex state of the tissue during gastrulation.

A similar mechanism of stabilization by Scab-dependent friction has been discovered in beetle embryos, but in a very different developmental context and mediated by a different integrin (Münster et al., 2019). Nevertheless, this suggests that the process is either conserved or has been co-opted in *Drosophila*. On the other hand, in some Dipteran species (Lemke and Schmidt-Ott, 2009), germ-band twists occur seemingly unchecked. It will be interesting to examine whether the Scab-dependent friction mechanism is at play in other Dipteran species and whether germ-band destabilization by twisting incurs a fitness cost for affected embryos.

Our observation that the twist of the germ band is biased to the left suggests that an intrinsic L/R asymmetry of the cells contributes to germ band instability. In *Drosophila*, L/R asymmetric tissue-level processes originate from the intrinsic asymmetry of epithelial cells, driven by the molecular motor Myo1D (Ishibashi et al., 2020; Sato et al., 2015; Taniguchi et al., 2011). Since ectopic Myo1D expression is sufficient to induce *de novo* chirality in many tissues (Lebreton et al., 2018; Petzoldt et al., 2012b; Spéder et al., 2006), it was reasonable to hypothesize that Myo1D expression during the most unstable period of germ band extension would affect the chiral properties of that tissue. Our functional experiments are consistent with this hypothesis. We have shown that silencing zygotic Myo1D expression removes the twisting bias and that Myo1D overexpression induces a substantial increase in the percentage of embryos displaying leftward twisting of the germ band during extension. We therefore propose that the intrinsic chirality of the epithelial tissue could be a factor in germ band extension instability. Unlike other chiral processes in *Drosophila*, the leftward bias of the germ band twist is less pronounced. This could indicate that the twisting is indeed an emergent mechanical phenomenon, consistent with our model, rather than a functionally required asymmetry. Similarly, the mutation of the *rotated abdomen* locus induces altered laterality in a series of chiral organs in the adult fly, but also a half-body rotation in the larva, revealing a latent tissue chirality present at that stage (Martín-Blanco and García-Bellido, 1996). Since the zygotic expression of Myo1D is destined for the hindgut, where the chiral phenotype manifests fully, it may not have a specific function during the germ band extension. Rather, the instability is a nuisance that the developmental system must address to ensure stable germ band progression. Another obvious solution to this problem that evolution may have discovered would be to suppress early Myo1D expression. Comparative studies of *myo1D* and *scab* in relation to the twisting germ band phenotype across Dipteran phylogeny may shed light on the evolution of compensatory mechanisms that ensure robustness in deterministic developmental systems under mechanical stress.

## Materials and Methods

### Fly stocks and genetics

The *D. melanogaster* stocks utilized in this study are listed in Suppl. Table XI. Most of the stocks were obtained from public collections (Bloomington Drosophila Stock Center, BDSC or Kyoto Stock Center, KSC) as indicated. The fly lines of the necessary genotype have been produced by genetic crosses following standard procedures. To identify embryos homozygous for the various mutant alleles, lines carrying balancers expressing fluorescent proteins in embryos were established. The fluorescent balancers utilized were *CyO, twi-GFP* (gift from Akanksha Jain) and *TM3, Kr-GFP* (BDSC, FBst0005195). The fluorescent membrane marker *Gap43::mCherry* is a gift from Kassiani Skouloudaki (FBal0258719). The construct has been recombined to obtain the homozygous viable line *w*;;RGAP43::mCherry*. The line *w*; scb^2^/ CyO, twi-GFP; RGAP43::mCherry* has been generated to study the effect of *scab* loss of function allele in early embryogenesis in live imaging experiments. The constructs *Myo1D^K2^*, *UAS-Myo1D::GFP* and *Myo1D::mNeonGreen* are a gift from Stéphane Noselli (Lebreton et al., 2018). The line *UAS-Myo1D::GFP; da-Gal4* has been generated for this study. The *Scab::mNeonGreen* reporter construct has been produced in collaboration with the Genomic Engineering Facility of MPI-CBG according to the FlyFos protocol (Sarov et al., 2016). The reporter lines *w*; shg-mCherry; Scab::mNeonGreen* and *w; Myo1D::mNeonGreen; RGAP43::mCherry* have been generated for this study. Embryos of the genotypes *da-Gal4; UAS-Myo1D-RNAi* and *Myo1D::mCherry; Scab::mNeonGreen* were generated by crossing flies.

### Animal husbandry and embryo collection

To collect embryos of the desired developmental stage, virgins and males of the appropriate genotype were caged in a ratio 1:3 in presence of freshly-prepared yeast paste (dry yeast and water) and let adjust to the cage for a minimum of 3 days. Before the egglays, a pre-lay of at least 1h 30’ was used to ensure the synchronization of the collected embryos. The egglays were performed in a time window spanning from 12:00 pm to 3 pm, on agarose-apple juice plates in presence of fresh yeast paste. The embryos were collected with a brush from the agar plate and transferred to a cell strainer with 100 μm nylon mesh for dechorionation.

### Production of the Scab::mNG fly line

The Scab::mNG fly line has been produced according to the published FlyFos protocol (Sarov et al., 2016). The FlyFos030776 Fosmid has been modified by recombineering with the insertion of the membrane-bound mNeonGreen (mNG; FPbase ID ZRKRV) coding sequences at the C-terminus of the *scab* coding sequence, replacing the sGFP tag. The DNA construct was injected in embryos from the landing line VK00033 to obtain the insertion of the transgene on the third chromosome (Venken et al., 2006). The recombinant flies have been selected using the dsRed fluorescent marker in the eyes and the developing brain (Sarov et al., 2016), and the balanced fly stocks have been established through the standard sequence of crosses.

### Sample mounting and imaging techniques

#### Live embryos

The embryos were dechorionated in a 20% solution of NaClO in ddH_2_O for 90 sec, thoroughly rinsed with water, then glued on the outside of glass capillaries or slides with heptane glue (Dahmann, 2016).

##### Brightfield imaging

Dechorionated embryos were transferred to a 24×60 mm coverslip coated with a thin layer of heptane glue. To prevent desiccation during the imaging procedure, the embryos were covered with a drop of Halocarbon Oil 700 (Sigma-Aldrich). The samples were then imaged by wide-field Differential Interference Contrast (DIC) and epifluorescence microscopy with an upright Axioplan 2 Fluorescence Phase contrast Microscope (Zeiss) equipped with Plan-NEOFLUAR, DIC, AxioCam HRc. The imaging has been conducted at room temperature. A single focal plane was acquired for each embryo every 2 min using the 10x/0.30 Plan-NEOFLUAR air objective (Zeiss) with the ZEN pro 3.8 software (Zeiss). The movies were processed in Fiji (Schindelin et al., 2012) as described in “Processing and analysis of images”.

##### Laser-scanning confocal microscopy

Dechorionated embryos were glued at the bottom of a MatTeK dish with a thin layer of heptane glue and imaged in water immersion. The samples were imaged by laser-scanning confocal microscopy with an upright Zeiss LSM 880 Airy upright microscope, equipped with two PMT and one GaAsP detector, using the LD LCI Plan Apo 25x/0.8 immersion objective (Zeiss) in Water immersion. Serial sections with an optimal Z-resolution step of 1.24 μm were acquired via Spectral Acquisition with ZEN 2.3 SP1 FP3 (black version). The images were processed in Fiji (Schindelin et al., 2012) as described in “Processing and analysis of images”.

##### Light sheet imaging

Several rows (4 to 5) of up to 20 dechorionated embryos were aligned on an agar pad with a fine plastic hair paintbrush and oriented with the imaged side facing the agar. Each line of embryos was then transferred to a pre-coated capillary (Z328472 or Z328480, Sigma) covered with heptane glue, by bringing the capillary in contact with the embryos. Red capillaries could hold 4 lines of embryos, at 90°C of each other, with the potential of correction of the orientation within the microscope, while the black capillaries could hold up to 5 lines of embryos. The samples were then imaged with Zeiss Light sheet Z.1 or Zeiss Light sheet7 using the Zeiss Plan Apo 20x/1.0 Water DIC objective or 40X (Fig. 2C). The imaging has been conducted at a constant temperature of 25°C unless specified otherwise. For the *w*;;RGAP43::mCherry* and *w*; scb^2^/ CyO, twi-GFP; RGAP43::mCherry* lines, z-stacks with a spacing of 2 μm and covering around 60% of the volume of the embryo were acquired every 2 min with ZEN 2014 SP1 9.2.10.54 or ZEN 3.1 9.3.7.393 software (Zeiss). The movies were processed in Fiji (Schindelin et al., 2012) as described in “Processing and analysis of images”.

For the *w*; shg-mCherry; Scab::mNeonGreen* line, 3 color imaging was performed at 18°C, in PBS with optimal laser power to detect early signal of E-cadherin and Scab while preventing photobleaching over time. A 561 nm laser with laser power 8%, time exposure 25 ms was used in combination with 575-615 nm filter, a 488 nm laser at 10% for 30 ms with 505-545 nm filter and the autofluorescence of yolk and vitelline membrane was detected with the 405 nm laser with 3% laser power and 10 ms exposure with the same filter. This last channel was used to remove the autofluorescence signal detected by the 488 nm excitation. Z-stack spacing was consistent per dataset and could vary between 0.7 μm up to 1.5 μm, necessary for good Local Projection (see “Processing and analysis of images”). The *w; Myo1D::mNeonGreen; RGAP43::mCherry* line was imaged at 18°C, in PBS with the same parameters and the z-spacing was 1µm at 18°C.

For the tissue flow analysis, multi-view light-sheet imaging was performed on a Zeiss Light sheet 7 microscope equipped with a 20 x/1NA Plan-Apochromat water immersion objective to acquire stacks with 0.28 μm XY-resolution and 2 μm Z-step covering two thirds of the volume of the embryo in a single view. RGAP43::mCherry embryos were embedded in glass capillaries in 2% low melting agarose dissolved in PBS, oriented with the lateral side towards the objective and kept at 25° C. Excitation and collection of mCherry was achieved with a 561 nm laser and LP560 dichromatic mirror. Two angles (left and right side) were acquired every 30 seconds and posteriorly registered using the Multiview reconstruction plugin in Fiji (Heemskerk and Streichan, 2015; Schmied et al., 2016).

#### Fixed embryos

The embryos were dechorionated in a 50% solution of NaClO in ddH_2_O for 2 min, thoroughly rinsed with water and fixed in 4% PFA in PBS according to the published protocol (Tomancak et al., 2002). The samples were stored in 100% MeOH at -20 C in the dark.

##### Brightfield microscopy

after the staining procedure, the embryos were mounted in 80% Glycerol solution in 1× PBS on glass slides with 24×60 mm coverslips and glass spacers. The samples were then imaged by wide-field Differential Interference Contrast (DIC) microscopy with an upright Apotome (Zeiss Axio Imager.Z2), using the 10x/0.45 and 20x/0.45 air objectives (Zeiss). Serial sections were acquired every 3 µm with ZEN pro 3.8 software (Zeiss). The images were processed through the Extended Depth of Field function of the software.

##### Laser-scanning confocal microscopy

after the staining procedure (see “RNA in situ hybridization techniques”), the embryos were mounted in SlowFade Diamond mounting medium (Thermo Fisher Scientific). The samples were placed between a 24×50 and a 22×22 mm coverslip using three layers of transparent tape as spacers, so that both sides of the samples could be imaged. The samples were imaged by laser-scanning confocal microscopy (Zeiss LSM 700 inverted or Zeiss LSM 880 Airy upright, equipped with two PMTs and one GaAsP detector) using respectively the Zeiss Plan Apo 25x/0.8 or the LD LCI Plan Apo 25x/0.8 immersion objectives (Zeiss) in Glycerol immersion. Serial sections with a z-resolution of 2 μm and covering around 60% of the embryo’s volume were acquired with Zeiss’s ZEN 2012 SP5 FP3 (black version) or ZEN 2.3 SP1 FP3 (black version). The images were processed in Fiji (Schindelin et al., 2012) to obtain the Maximum Intensity Projections.

##### Electron Microscopy

*Oregon R* or *RGAP43::mCherry* embryos were dechorionated with 50% bleach for 2min, staged on agar, transferred with a paintbrush to a 150 µm deep Cu carrier filled with 20% BSA, then covered by the 1-Hexadecene coated flat side of a 300 µm carrier which was pre-wetted to prevent air bubble trapping when closing. After high pressure freezing in a Leica ICE high-pressure freezer, the carriers were transferred to a freeze substitution cocktail of either 0.1% UA, 1% Os_2_O_4_, 0,5% GA, 4% H_2_0 in acetone or 1% OsO_4_, 0.5% UA, 3% H_2_O in Acetone – and slowly brought to 0° Cover days. The resin embedding was performed in Araldite or Epon Hard with decreasing concentrations of acetone over several days. The resin was polymerized at 60°C for at least 3 days. The embryos were oriented laterally for the flat embedding and ultra-sectioned (70 nm) at different depths with a Leica EM UC6 ultramicrotomes, on formvar coated slot grids and at stages when *scab* was expressed. Post-staining was performed with 1% or 2% UA and Lead Citrate for varying times. Images were acquired with a FEI Tecnai 12 TEM (100kV) and a TVIPS F416 4kx4k CCD camera. Sections shown have more or less contrast on the membrane and often have ice contamination, because of the integrity of the vitelline membrane.

### RNA in situ hybridization techniques

The experiments were performed according to the Berkeley Drosophila Genome Project RNA *in situ* hybridization protocol (Tomancak et al., 2002) or an optimized version of the published Hybridization Chain Reaction (HCR) RNA-FISH protocol available from Molecular Instruments, in the version for whole-mount *Drosophila* embryo samples (Choi et al., 2018). The following adaptations were implemented: the Protease K incubation was 7 min at room temperature with 0.4 μL of stock solution (20 mg/mL) in 2 mL PBT 0.1% and a 20 min long post-fixation was performed in 4% formaldehyde in PBT 0.1%, rinsed 6 times with PBT 0.1%.

### Processing and analysis of images

#### Quantification of germ band deformation

##### Tortuosity

The tortuosity measurements have been performed in ImageJ Fiji according to the following steps:

1) Selection from the live recording data of the frames corresponding to the three timepoints (Tps) of interest (Tp 1 = the moment before the closure of the PMG invagination when the pouch assumes a triangular shape; Tp 3 = the moment when germ band extension stops; Tp 2 = the moment when the germ band reaches half of its full extension, defined at Tp 3)
2) Rendering of the frames in 3D via the 3D Script Fiji plugin (Schmid et al., 2019).
3) Definition of the ROI by drawing a segmented line over the midline of the germ band, from the emergence of the ventral furrow at the posterior end of the embryo until the tip of the germ band.
4) For each ROI, measurement of the length and the Feret Diameter, corresponding to the Euclidean distance between the two extremities of the ROI. From these two parameters the tortuosity of the line has been calculated as the ratio of length to Feret Diameter

##### Deviation from straightness

This method was developed to quantify germ band deformation in 3D, capturing its deviation from the typical straight alignment along the longitudinal axis of the embryo in a more comprehensive manner. The position of multiple interest points along the mid-line of the germ band has been placed in the 3D dataset with the help of the Mastodon plugin in ImageJ Fiji (Girstmair et al., 2025). The two points marking the extremities of the germ band midline, called “germ band tip” (GB tip) and “ventral furrow point” (VF point) have been identified and tracked over time. The distance of each point of interest from a plane passing through the center of the embryo along its longitudinal axis has then been calculated, as explained in Supplementary Methods.

#### Live recordings of Scab and Myo1D reporter expression

To distinguish the signal associated with mNeonGreen from the autofluorescence of the vitelline membrane and the yolk, also excited by the 488 nm laser and detected with a GFP filter, the imaging channel from the 405 nm excitation was used as a mask to clear out the autofluorescence from the 488 nm channel. This was necessary as the weak dorsal localization of Scab and Myo1D are in close proximity to the vitelline membrane and superimposed with the yolk in projections. A script was used to first apply a median filter with a radius of 0.8 px, then a Gaussian blur of 1.2 px. Different background subtractions were used for each channel: notably, a radius of 50 px for the mNG channel. An extra Gaussian filter was applied to the 405 nm channel, which was then thresholded with a constant intensity threshold, and the created mask was used to clear the corresponding signal in every slice of the 488 nm stack. The same process was applied to the Ecad::mCherry channel. This allowed for a local z-projection, with the reference surface calculated from the membrane labelled channel (Ecad::mCherry or RGAP::43mCherry) (Herbert et al., 2021), which revealed the localization of the apical dorsal recruitment of Scab and Myo1D.

Scab intensity was measured in an adjusted ROI by creating a mask based on the intensity thresholding of the highest signal level on a Gaussian-blurred image with a radius of 8 px. Additionally, an area-based exclusion filter was used to only keep the largest thresholded area for the measurement. This mask was applied to the original local z-projection with the Scab-mNG data and the intensity measurement was performed in this ROI. This helped refine the intensity variations as well as giving information about the area expressing Scab. For Myo1D quantification, the intensity was measured in the entirety of the local z-projection in the corresponding channel.

As multi-color light sheet imaging requires different manual alignment offsets for each laser, and as the quality of the alignment is judged on features of the samples like the vitelline envelope, the slightest mismatch between the 405 nm channel and the others can require an extra dilatation of the mask to ensure that the autofluorescence signal from the vitelline envelope is completely removed. This process can result in an artifact suggesting that some areas are depleted of the protein of interest. While the dataset Fig. 2 suggests that there is no Scab along the border of the amnioserosa, Fig. S3 shows through the stack slices that Scab is present in the area.

For the quantification of the apical signal of Scab as a heat map (Fig. 2E), a moving ROI following the posterior midgut (area expressing scab) was defined on a stack of a dorsally mounted embryo. The integrated intensity was measured in this ROI for each slice (over 13 slices) and over time.

#### Quantification of HCR RNA-FISH

The signal intensity has been quantified in ImageJ Fiji on the maximum projection of the images, without background subtraction. The acquisition settings were kept the same for every replicate, the laser power and gain values were the same between samples of the same replicate (control and experiment). The contrast was set to the same values between the control and experiment of the same replicate. The intensity was quantified over a 10-pixel-thick segmented line drawn on the area of *myo1D* expression domain. The signal reduction was quantified relative to the positive control (*Oregon R* embryos expressing *myo1D* normally).

#### Tissue flow analysis

##### Post-processing

the 2D + time cartographic projections were analyzed with the iterativePIV plugin by Qingzong Tseng (Bailles et al., 2019; Münster et al., 2019; Tseng et al., 2012). For each time point, displacement fields were computed and exported for post-processing in a custom Jupyter notebook pipeline, where strain (*E*) and curl (*R*) rates were calculated as

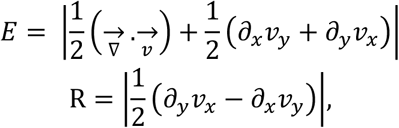

where *v* corresponds to the displacement obtained from PIV divided by the time interval between frames (minutes). These values were used to generate the color-coded overlays shown in Fig. 4.

To generate the temporal line plots, tissue flow fields were analyzed to identify the position and trajectory of the center of rotation. Ten sampling points were defined along the axis of rotational displacement surrounding this region, capturing the dominant direction of tissue deformation. Strain and curl rates were extracted at these positions for each time point and averaged to obtain a representative temporal profile of tissue dynamics.

To quantify left–right asymmetry, measurements were grouped according to their position relative to the PMG midline and averaged separately for the left and right sides. The asymmetry index was calculated as

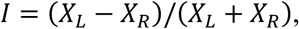

where *X* represents the mean magnitude of either curl or strain rates on the left (L) or right (R) sides. This metric provides a normalized, dimensionless measure of asymmetry, where positive values indicate higher curl on the left side, negative values indicate higher curl on the right side, and zero represents perfect bilateral symmetry.

##### Quantification of cellular features at gastrulation

using the cartographic projections, two square regions of interest (ROIs; 360 × 360 pixels) were defined on each side of the PMG. Cells within each ROI were segmented, labeled, and fitted with ellipses using the MorphoLibJ plugin (Legland et al., 2016; “MorphoLibJ,” 2026), and morphological features were extracted for each cell. The resulting measurements were exported and analyzed in a Jupyter notebook pipeline. Cell orientation was redefined such that 0° corresponded to the anteroposterior (AP) axis. Orientations from the left and right sides were compared using absolute values to account for mirrored geometry across the midline. Mean cell orientation and cell elongation (minor-to-major axis ratio) were computed over time to generate the line plots shown in Fig. 4.

### Laterality scoring

The handedness of the germ band twist in fixed embryos showing altered germ band extension phenotype has been manually scored by observing them one-by-one at the stereomicroscope (brightfield or epifluorescence). The handedness of *in vivo* data has been scored from the acquired 3D time-lapse recording. The relative position of the tip of the germ band to the longitudinal axis of the embryo has been used to define the direction of the twist (tip to the Left or tip to the Right side of the embryo). For the scoring, all the samples were observed from the dorsal side, oriented as shown in Fig. 4A. In the fixed samples analysis, embryos ranging from stage 7 to stage 11 of germ band extension have been scored. Embryos showing further morphological alterations in addition to the curvature of the germ band, as well as embryos in which the position of the head was rotated, but the shape of the germ band remained unaffected, were excluded from the scoring, since the morphological alterations could not be unambiguously attributed to germ band instability. For the analysis of fixed samples in the *myo1D* knockdown or Myo1D::GFP overexpression experiments the samples have been randomized with negative controls before scoring, and the scoring has been performed blind. Multiple replicates have been scored for each genotype, and each replicate consists of two embryo collections, 2-5 and 5-7 hours after egg laying. Embryos of the desired stages have been selected manually. Every imaging set-up has its own optical paths that could lead to the mirroring of the data. Therefore, attention was paid to ensure the proper attribution of the left and right directions across all the data collected.

### Myo1D downregulation and overexpression experiments

To obtain downregulation of zygotic *myo1D* expression via RNA interference, virgins expressing the *da-Gal4* ubiquitous maternal driver have been crossed with males homozygous for a *UAS-myo1D-RNAi* construct (line #33971 from BDSC) or a *UAS-GFP-RNAi* construct (line #56181 from BDSC) as a negative control. To obtain the maternal ubiquitous overexpression of Myo1D::GFP, driver and responder constructs have been combined in the new line *w*; UAS-Myo1D::GFP; da-Gal4*.

### Statistical analysis

The statistical data analyses were performed using Rstudio version 4.3.2. The level of significance was set at p < 0.05. The specific statistical method applied to each experiment are indicated in the Figure captions and in the relative section of the Results, as well as the sample sizes. Further details about each analysis are provided in the related Supplementary Tables.

For the analysis of germ band deformation in 3D, a one-way Mann Whitney test has been performed, testing *scab* mutant greater than wt. To account for repeated measurements, the Bonferroni Correction has been applied to the p-values, as indicated in the Suppl. Tables.

The analysis of the binary developmental outcomes (twist penetrance [1/0] and twist bias [right/left]) has been conducted employing generalized linear mixed-effects models (GLMMs) with a binomial family distribution. The models were fitted to the data using the glmer() function from the “lme4” package. In these models, the experimental condition was specified as a fixed effect, while the biological replicate was included as a random effect to account for batch-to-batch variability. The model estimates (log-odds) were exponentiated to calculate Odds Ratios, which served as the measure for the magnitude of the effect.

### Mathematical model

The construction of the mathematical model is detailed in the Supplemental Note about the model, which includes Refs. (Abramowitz and Stegun, 1964; Carmo, 1976; Cholakova et al., 2021; De Canio et al., 2017; Fily et al., 2020; Lauga, 2020; Man and Kanso, 2019; Zhao and Haas, 2025).

## Acknowledgements

We thank current and former members of the Tomančák (LoPaTs) and Haas groups for discussions and support throughout this project; Stéphane Noselli for sharing the Myo1D-related fly lines; Arthur Boutillon for performing the test experiment of Myo1D::mNG live spectral acquisition; Anaïs Bailles and Bipasha Dey for critically reading the manuscript; the Light Microscopy Facility of the Max Planck Institute of Molecular Cell Biology and Genetics (MPI-CBG) for assistance with data acquisition; Michaela Yuan, Weihua Lang, and Tobias Fürstenhaupt for assistance with electron microscopy; the Genomic engineering Facility of MPI-CBG for assistance with the Flyfos protocol; Sven Ssykor and Cornelia Maas for flykeeping; Federica Luppino, Pierre Mangeol, and Rubén González Miguélez for consultations and help with the statistical analyses; and Nuno Pimpão Martins for suggesting the tortuosity measurement.

## Authors contributions

GS and PT conceived the study. GS designed experiments, generated fly stocks, acquired live and fixed microscopy data, performed in situ hybridization, processed and analyzed the live germ band extension data, performed the tortuosity measurements and the twist penetrance and handedness scorings. MBC acquired the multiview data sets, conceived and conducted the tissue flow analysis and performed the cellular level quantifications. MS and PAH derived and analyzed the theoretical models. GS and CB generated the transgenic Scab::mNG reporter. CB generated fly stocks, prepared and imaged the samples for electron microscopy and performed the imaging, processing and analyses of the Scab and Myo1D reporter line expressions. MA created the computational tools for the germ band deviation analysis. PM supported with the fly rearing and performed some of the *in-situ* hybridization staining. GS wrote the initial version of the manuscript. All the authors revised and contributed to the text. GS, PAH and PT polished the final version of the manuscript.

