## Supplementary Tables for "“Embryo-eggshell interaction counteracts chiral bias in early *Drosophila* morphogenesis”"

**Suppl. Table I** = penetrance of the twisting phenotype in wild-type and *scab* mutant embryos (p-values; Fisher's exact test, one tail)

| Genotype | p-value | p < 0.05 |
|---|---|---|
| Oregon R – $w^{1118}$ | 0.97 | |
| $w^{1118}$ – RGAP43::mCherry | 0.45 | |
| Oregon R – RGAP::mCherry | 0.95 | |
| RGAP43::mCherry – $scb^2/scb^2$ | 0.00016 | Y |

Sample sizes: Oregon R (n = 55); $w^{1118}$ (n = 44); RGAP43::mCherry (n = 71); $scb^2/scb^2$ (n = 21)

**Suppl. Table II** = comparison of germ band tortuosity in wild-type (*RGAP::mCherry/RGAP::mCherry*; n = 61) and *scab*$^{-/-}$ mutant embryos (n = 19) at the three timepoints of interest (p-values; Mann Whitney test, one way testing *scab* mutant greater than wt).

| Timepoint | p-value | p < 0.05 |
|---|---|---|
| 1 | $7.9\times10^{-5}$ | Y |
| 2 | 0.0013 | Y |
| 3 | 0.000101 | Y |

**Suppl. Table III** = comparison of the germ band's deviation from straightness in wild-type (*RGAP::mCherry/RGAP::mCherry*; n = 66) and *scab*$^{-/-}$ mutant embryos (n = 21) at the 15 timepoints of interest (p-values; Mann Whitney test, one way testing *scab* mutant greater than wt)

| **Timepoint** | **p-value** | **p < 0.0033 (Bonf. Corr.)** |
|---|---|---|
| 0 | 0.21 | |
| 1 | 0.024 | |
| 2 | 0.013 | |
| 3 | 0.0064 | |
| 4 | 0.052 | |
| 5 | 0.0074 | |
| 6 | 0.0103 | |
| 7 | 0.0004 | Y |
| 8 | 0.0028 | Y |
| 9 | 0.0044 | |
| 10 | 0.0013 | Y |
| 11 | 0.0016 | Y |
| 12 | 0.0027 | Y |
| 13 | 0.0017 | Y |
| 14 | 0.0038 | |

Note: the Bonferroni Correction (Bonf. Corr.) has been applied to account for repeated measurements.

**Suppl. Table IV** = comparison of deviation from straightness of the germ band tip in wild-type (*RGAP::mCherry/RGAP::mCherry*; n = 66) and *scab*$^{-/-}$ mutant embryos (n = 21) at the 15 timepoints of interest (p-values; Mann Whitney test, one way testing *scab* mutant greater than wt).

| **Timepoint** | **p-value** | **p < 0.0033 (Bonf. Corr.)** |
|---|---|---|
| 0 | 0.11 | |
| 1 | 0.052 | |
| 2 | 0.014 | |
| 3 | 0.00702 | |
| 4 | 0.13 | |
| 5 | 0.0032 | Y |
| 6 | 0.075 | |
| 7 | 0.00096 | Y |
| 8 | 0.0056 | |
| 9 | 0.00075 | Y |
| 10 | 0.00096 | Y |
| 11 | 0.0012 | Y |
| 12 | 0.0023 | Y |
| 13 | 0.00099 | Y |
| 14 | 0.00087 | Y |

Note: the Bonferroni Correction (Bonf. Corr.) has been applied to account for repeated measurements.

**Suppl. Table V** = comparison of deviation from straightness of the ventral furrow point in wild-type (*RGAP::mCherry/RGAP::mCherry*; n = 66) and *scab*$^{-/-}$ mutant embryos (n = 21) at the 15 timepoints of interest (p-values; Mann Whitney test, one way testing *scab* mutant greater than wt)

| **Timepoint** | **p-value** | **p < 0.0033 (Bonf. Corr.)** |
|---|---|---|
| 0 | 0.605 | |
| 1 | 0.0103 | |
| 2 | 0.012 | |
| 3 | 0.14 | |
| 4 | 0.21 | |
| 5 | 0.17 | |
| 6 | 0.028 | |
| 7 | 0.016 | |
| 8 | 0.43 | |
| 9 | 0.18 | |
| 10 | 0.60 | |
| 11 | 0.62 | |
| 12 | 0.68 | |
| 13 | 0.63 | |
| 14 | 0.74 | |

Note: the Bonferroni Correction (Bonf. Corr.) has been applied to account for repeated measurements.

**Suppl. Table VI** = significance of the twist handedness bias in different destabilizing genetic backgrounds (p-values; Binomial test)

| Type of experiment | Stock/genotype | n | p-value | $p < 0.05$ |
|---|---|---|---|---|
| **Fixed imaging data** | BDSC # 3098 | 27 | 0.13 | |
| | Δ7675-4 line | 119 | $1.5\times10^{-7}$ | Y |
| | Δ7675-3 line | 160 | $2.3\times10^{-5}$ | Y |
| | Δ7683-4 line | 209 | $3.7\times10^{-6}$ | Y |
| | Δ7683-5 line | 124 | $7.4\times10^{-9}$ | Y |
| | Δ7675-2 line | 99 | 0.0055 | Y |
| | Δ7675-1 line | 148 | $1.7\times10^{-8}$ | Y |
| | Δ7730-3 line | 98 | $8.1\times10^{-6}$ | Y |
| | Δ7730-2 line | 85 | $1.1\times10^{-6}$ | Y |
| | Δ7730-4 line | 170 | $5.6\times10^{-8}$ | Y |
| | Δ7683-2 line | 107 | 0.00012 | Y |
| | Δ7683-3 line | 172 | 0.00090 | Y |
| | Δ7624-1 line | 111 | 0.035 | Y |
| | Δ7683-1 line | 100 | $6.3\times10^{-8}$ | Y |
| | Δ7675-5 line | 35 | 0.00019 | Y |
| | Δ7730-1 line | 26 | 0.14 | Y |
| **BF live imaging data** | *Oregon R* | 29 | 0.064 | Y |
| | $w^{1118}$ | 16 | 0.028 | Y |
| | Δ7675/Δ7675 | 13 | 0.16 | |
| | Δ7675-1/Δ7675-1 | 8 | 0.11 | |
| | Δ7675-3/Δ7675-3 | 44 | 0.039 | Y |
| | Δ7675-4/Δ7675-4 | 27 | 0.063 | |

**Suppl. Table VII** = comparison of tissue dynamics and cellular features on the two sides of the posterior-midgut invagination in gastrulating *RGAP::mCherry* embryos, n = 10 (p-values; Wilcoxon signed-rank test, one sided)

| **Type of analysis** | **Timepoint** | **p-value** | **p < 0.05** |
|---|---|---|---|
| Curl rate | 15 min after cellularization | 0.00098 | Y |
| Strain rate | 15 min after cellularization | 0.00098 | Y |
| Ratio minor/major axis versus time | 15 min after cellularization | 0.049 | Y |

Note: curl and strain rate have been analyzed on a sample group of 10 embryos. The ratio of minor to major axis against time has been analyzed on a different sample.

**Suppl. Table VIII** = significance of the twist handedness bias in samples with altered Myo1D expression (p-values, Binomial test)

| Type of experiment | Genotype | n | p-value | $p < 0.05$ |
|---|---|---|---|---|
| **Downregulation** | Oregon R _repl 1 | 85 | 0.023 | Y |
| | Oregon R _repl 2 | 142 | 0.0088 | Y |
| | Oregon R _repl 3 | 37 | 0.068 | Y |
| | da>myo1D-RNAi _repl 1 | 26 | 0.14 | |
| | da>myo1D-RNAi _repl 2 | 78 | 0.081 | |
| | da>myo1D-RNAi _repl 3 | 85 | 0.054 | |
| | da>GFP-RNAi _repl 1 | 52 | 0.0049 | Y |
| | da>GFP-RNAi _repl 2 | 99 | 0.0011 | Y |
| **Maternal Overexpression** | Oregon R _repl 1 | 262 | $7.8{\times}10^{-5}$ | Y |
| | Oregon R _repl 2 | 362 | 0.00020 | Y |
| | da>Myo1D::GFP _repl 1 | 224 | $5.4{\times}10^{-12}$ | Y |
| | da>Myo1D::GFP _repl 2 | 335 | $4.0{\times}10^{-13}$ | Y |

**Suppl. Table IX** = comparison of the twist handedness bias in samples with altered Myo1D expression [p-values and odds ratio percentage (magnitude); Binomial Generalized Linear Models (GLMMs)]

| Experiment | Genotype | p-value | p< 0.05 | Odds Ratio Percentage |
|---|---|---|---|---|
| **Downregulation** | wt – da>GFP-RNAi | 0.17 | | 0.747 (CI 0.493 – 1.132) |
| | wt – da>Myo1D-RNAi | $2.6\times10^{-2}$ | Y | 1.531 (CI 1.051 – 2.231) |
| **Maternal Overexpression** | wt – da>Myo1D::GFP | $9.9\times10^{-5}$ | Y | 0.618 (CI 0.485 – 0.788) |

Note: The OR represents the practical magnitude of the effect. An OR = 1 means no difference compared to the reference condition. An OR > 1 means the outcome is more likely to occur (e.g., an OR of 2.5 means the condition has 2.5 times the odds of the event occurring). An OR < 1 means the outcome is less likely to occur.

CI: "Confidence Interval"

**Suppl. Table X** = twist penetrance in samples with altered Myo1D expression [p-values, odds ratio percentage (magnitude); Binomial Generalized Linear Models (GLMMs)]

| Experiment | Genotype | p-value | p< 0.05 | Odds Ratio Percentage |
|---|---|---|---|---|
| **Downregulation** | wt – da>GFP-RNAi | $1.1\times10^{-3}$ | Y | 1.463 (CI 1.165 – 1.838) |
| | wt – da>myo1D-RNAi | $1.8\times10^{-6}$ | Y | 1.813 (CI 1.420 – 2.314) |
| **Maternal Overexpression** | wt – da>Myo1D::GFP | $3.0\times10^{-72}$ | Y | 4.415 (CI 3.775 – 5.190) |

Note: The OR represents the practical magnitude of the effect. An OR = 1 means no difference compared to the reference condition. An OR > 1 means the outcome is more likely to occur (e.g., an OR of 2.5 means the condition has 2.5 times the odds of the event occurring). An OR < 1 means the outcome is less likely to occur.

CI: "Confidence Interval"

**Suppl. Table XI** = signal reduction in samples undergoing *myo1D* RNAi knockdown (*da>myo1D-RNAi* embryos compared with *Oregon R* embryos (p-values; two-sample one-tailed T-test assuming unequal variances).

| Experiment | Genotype | p-value | p< 0.05 |
|---|---|---|---|
| **Stage 9** | Replicate 1 | $7.1\times10^{-6}$ | Y |
| | Replicate 2 | $3.1\times10^{-8}$ | Y |
| **Stage 10** | Replicate 1 | 0.0041 | Y |

**Suppl. Table XII** = genotypes of the fly strains used

| Genotype | ID | Source | Referred in the text asto as |
|---|---|---|---|
| *Oregon R* | - | MPI-CBG | |
| *w*$^{1118}$ | - | MPI-CBG | |
| *w*$^{*}$*;; sqh-Gal4,GAP43::mCherry / TM3, Sb*$^{1}$ | - | Tomancak Group | |
| *w*$^{*}$*;;RGAP43::mCherry* | - | this study | |
| *; cn*$^{1}$ *scb*$^{2}$ *bw*$^{1}$ *speck*$^{1}$*/ CyO;* | 3098 | BDSC | |
| *; if / Cy, twi-GFP; MKRS / TM6B, Tb*$^{1}$ | - | Tomancak Group | |
| *w*$^{*}$*; cn*$^{1}$ *scb*$^{2}$ *bw*$^{1}$ *speck*$^{1}$*/ Cy, twi-GFP ; RGAP43::mCherry* | - | this study | *scab*$^{2}$*/Cy; RGAPmCherry* |
| *y*$^{1}$ *w*$^{*}$*;D*$^{*}$ *gl*$^{3}$*/TM3, P{GAL4-Kr.C}DC2, P{UAS-GFP.S65T}DC10, Sb*$^{1}$ | 5195 | BDSC | |
| *w*$^{1118}$*;; Df(3R)Exel6196, P{XP-U}Exel6196/ TM6B, Tb*$^{1}$ | 7675 | BDSC | *Δ7675* |
| *w*$^{1118}$*;; Df(3R)Exel6204, P{XP-U}Exel6204/ TM6B, Tb*$^{1}$ | 7683 | BDSC | *Δ7683* |
| *w*$^{1118}$*;; Df(3R)Exel6263, P{XP-U}Exel6263/ TM6B, Tb*$^{1}$ | 7730 | BDSC | *Δ7730* |
| *w*$^{1118}$*;; Df(3R)Exel6145, P{XP-U}Exel6145/ TM6B, Tb*$^{1}$ | 7624 | BDSC | *Δ7624* |
| *w*$^{1118}$*;; Df(3R)Exel9014 / TM6B, Tb*$^{1}$ | 7992 | BDSC | *Δ7675-1* |
| *w*$^{1118}$*;; Df(3R)BSC489 / TM6C, Sb*$^{1}$ *cu*$^{1}$ | 24993 | BDSC | *Δ7675-2* |
| *w*$^{1118}$*;; Df(3R)ED6144 / TM6C, Sb*$^{1}$ | 150-339 | KSC | *Δ7675-3* |
| *w*$^{1118}$*;; Df(3R)ED6150 / TM6C, Sb*$^{1}$ | 150-340 | KSC | *Δ7675-4* |
| *w*$^{1118}$*;; Df(3R)ED6168 / TM6C, Sb*$^{1}$ | 150-342 | KSC | *Δ7675-5* |
| *w*$^{1118}$*;; Df(3R)BSC140 / TM6B, Tb*$^{1}$ | 9500 | BDSC | *Δ7683-1* |
| *;; Df(3R)Tl-P, e*$^{1}$ *ca*$^{1}$ */ TM3, Ser*$^{1}$ | 1910 | BDSC | *Δ7683-2* |
| *;; Df(3R)Tl-P, e*$^{1}$ *ca*$^{1}$*/ TM3, Ser*$^{1}$ | 106-725 | KSC | *Δ7683-3* |
| *w*$^{1118}$*;; Df(3R)BSC512 / TM6C, Sb*$^{1}$ *cu*$^{1}$ | 25016 | BDSC | *Δ7683-4* |
| *w*$^{1118}$*;; Df(3R)BSC752 / TM6C, Sb*$^{1}$ *cu*$^{1}$ | 26850 | BDSC | *Δ7683-5* |
| *w*$^{1118}$*;; Df(3R)BSC196 / TM6B, Tb*$^{1}$ | 9622 | BDSC | *Δ7730-1* |
| *w*$^{1118}$*;;Df(3R)ED5220, P{w*$^{+mW.Scer\backslash FRT.hs3}$*=3'.RS5+3.3'}ED5220 / TM3, Ser*$^{1}$ | 9200 | BDSC | *Δ7730-2* |
| *w*$^{1118}$*;; Df(3R)ED5220 / TM6C, Sb*$^{1}$ | 150-327 | KSC | *Δ7730-3* |
| *w*$^{1118}$*;; Df(3R)BSC222 / TM6B, Tb*$^{1}$ | 9699 | BDSC | *Δ7730-4* |
| *y*$^{1}$*, w*$^{*}$*, P{nos-phiC31int.NLS}X; PBac{y +-attP-3B}VK00033* | 32542 | BDSC | |
| *w*$^{*}$ *;; Scab::mNeonGreen* | - | this study | |
| *w*$^{*}$*; P{w*$^{+mW.hs}$*=FRT(w*$^{hs}$*)}G13 TI{TI}shg*$^{mCherry}$ | 59014 | BDSC | |
| *w*$^{*}$*; shg-mCherry; Scab::mNeonGreen* | - | this study | *Ecad::mCherry; Scab::mNeonGreen* |
| *;; da-Gal4 / da-Gal4* | 95282 | BDSC | |
| *w*$^{*}$*; P{UAS-rCD2.RFP.UAS-GFPi}attP40;* | 56181 | BDSC | |
| *y*$^{1}$ *sc*$^{*}$ *v*$^{1}$ *sev*$^{21}$*;; P{*$^{[+t7.7}$ *v*$^{+t1.8}$*=TRiP.HMS00928}attP2* | 33971 | BDSC | |
| *w; Myo1D*$^{K2}$ */ (CyO-Dfd-GFP); da-Gal4/ da-Gal4* | - | Noselli Group | |
| *w; UAS-chic-RNAi KK / CyO; UAS-Myo1D::GFP / (TM6Bf)* | - | Noselli Group | |
| *w; Myo1D::mNeonGreen / CyO;;* | - | Noselli Group | |

| | | |
|---|---|---|
| *w; Myo1D-mCherry / CyO;;* | - | Noselli Group |
| *w; Myo1D::mNeonGreen; RGAP43::mCherry* | - | this study |
