## Supplementary Methods for "“Embryo-eggshell interaction counteracts chiral bias in early *Drosophila* morphogenesis”"

<u>**Supplemental methods**</u>

### Germ Band Deviation Measurement

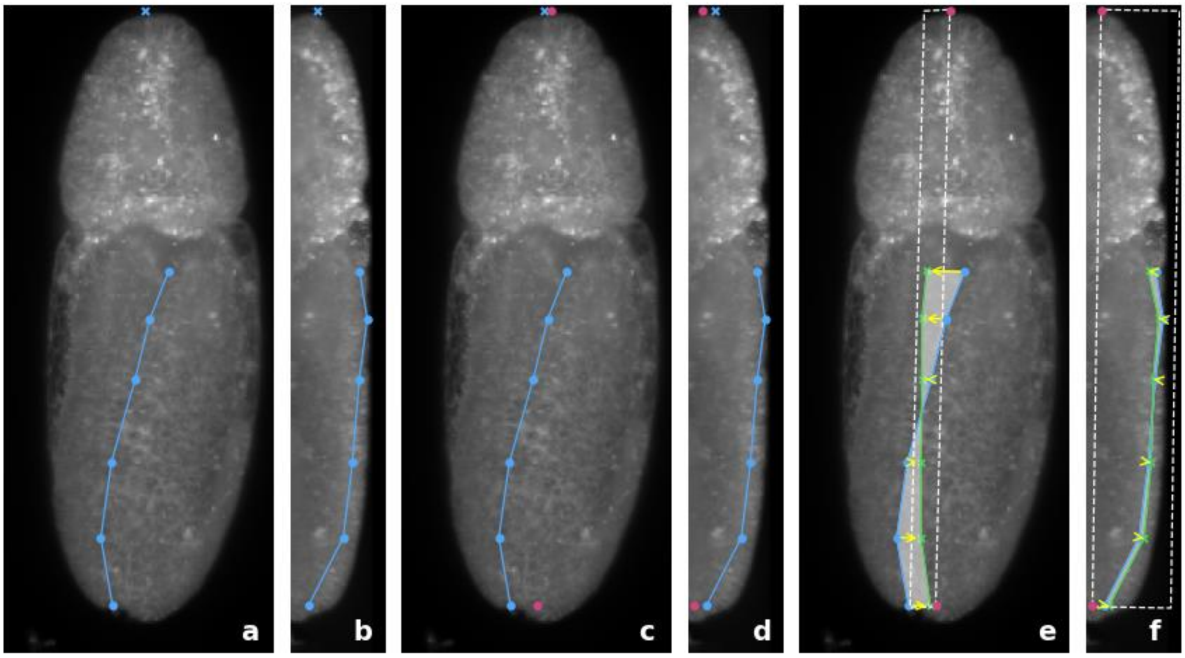


**Suppl. Methods Figure 1:** (a) Z-projection of a 3D image stack of a *Drosophila* embryo during germ band extension. The blue line and dots represent the manually traced germ band and the manually placed micropyle marker for the anterior end of the embryo. (b) similar to (a), but projected along the X-axis, providing a lateral view of the embryo. (c-d) shows the same features as (a) and (b), and in addition, the computed anchor points marking the anterior and posterior extremities intersecting the lateral symmetry 2D plane are highlighted in red. (e). The white dashed line represents an approximation of the lateral symmetry plane of the embryo. The blue line and dots depict the manually traced germ band, while the green line shows its projection onto the symmetry plane. Yellow arrows indicate the Euclidean distance between a point on the germ band and the approximated plane. (f) Same as (e), but projected along the X-axis.

In order to quantify the germ band deviation from the lateral symmetry plane in a 3D image stack capturing germ band extension in *Drosophila* embryos, we manually locate the germ band by placing points $v_1, \dots, v_n$ along its length, forming a polyline representation. Additionally, we manually locate the position of the micropyle and the posterior end of the embryo (Suppl. Methods Fig. 1a, b). The coordinates of all points are recorded as X, Y, Z positions in micrometers. A dedicated image analysis pipeline is used to segment the embryo and estimate the coordinates of the anterior and posterior anchor points for the midline 2D plane (see section “Approximation of the Anterior-Posterior Axis”).

We then approximate the lateral symmetry plane as the plane that contains the anterior and posterior anchors and minimizes the distance to the germ band (Suppl. Methods Fig. 1c–f). The germ band serves hereby as the best available landmark for determining the dorsal axis of the embryo.

Mathematically we compute the plane $P$ that contains the anterior-posterior anchors and minimizes the distance to the germ band. The distance $d(G, P)$ between the germ band $G$ and a plane $P$ is defined by

$$d(G,P) := \sqrt{\sum_{i=1}^{n} w_i \cdot d(v_i,P)^2}$$

where:

- $G := (v_1, v_2, \ldots, v_n)$ denotes the traced germ band.
- $w_i := \frac{1}{2L}\left(d(v_{i-1}, v_i) + d(v_i, v_{i+1})\right)$ denotes a weight corresponding to the point
- $L := \sum_{i=1}^{n-1} d(v_i, v_{i+1})$ is the length of the germ band
- $d(v_i, v_{i+1})$ denotes the Euclidean distance between the two points and
- $d(v_i, P)$ denotes the euclidean distance between a point and the plane

The minimal distance $d(G,P)$, attained by the optimal plane, defines the *germ band deviation*. This measure quantifies the average distance of the germ band to an $P$ optimally positioned lateral symmetry plane containing the anterior and posterior anchors. The unit for germ band deviation is µm.

**Ventral furrow deviation and GB tip deviation**

Beyond the global measurement, the same lateral symmetry plane can be used to assess the displacement of specific germ band landmarks. We measured the distance from the most posterior point of the germ band, termed the ventral furrow point (VF), and the most anterior point, referred to as the germ band tip (GB tip), to the symmetry plane (Suppl. Fig. S2I). These measurements define two additional metrics: *VF deviation* and *GB tip deviation* (Fig. 1G, H).

**Approximation of the Anterior-Posterior Axis**

We aim to determine the anterior-posterior axis of the embryos in our microscopy images. For a given embryo at a specific time point we aim to identify two reference points: an anterior and a posterior anchor within the 3D stack recorded by the microscope. The anterior anchor should be positioned at the center of the anterior tip of the embryo, while the posterior anchor is positioned at the posterior tip.

For this analysis, we use a downscaled version of the 3D image stack, with a pixel size of 2.3 µm and a slice spacing of 3 µm. The analysis also relies on the knowledge that the embryos were mounted in the microscope such that their anterior-posterior axis is approximately aligned with the Y-axis of the images. The image stack was then cropped to ensure the embryo is roughly centered in the frame.

The first step of our image analysis pipeline is to extract ten XZ slices of the 3D stack based on specific Y-coordinates as follows (Suppl. Methods Fig. 2a). Let $y_a$ and $y_b$ denote the Y-coordinates of the micropyle and the most posterior GB point, respectively (Suppl. Methods Fig. 1a, b). We generate slices for values of $y = (y_a - y_b)p + y_b$, where $p$ takes the values $\{0.05, 0.0625, 0.075, 0.0855, 0.1, 0.9, 0.9125, 0.925, 0.9375, 0.95\}$. At each of these Y-positions, a corresponding XZ slice is extracted.

For each of the ten slices (Suppl. Methods Fig. 2b), the following operations are performed: first, the intensity image is converted into a binary mask using Otsu thresholding (Suppl. Methods Fig. 2c). Holes in the mask are filled. Pixels located at the border of the mask are then extracted, and an ellipsoid is fitted to these border pixels (Suppl. Methods Fig. 2d). The center of the fitted ellipsoid is determined and transformed back into coordinates in the 3D image stack (Suppl. Methods Fig. 2e).

This procedure results in ten center coordinates corresponding to the ellipse centers (Suppl. Methods Fig. 2f, g). A line is then fitted through these center points, providing an estimate of the anterior-posterior axis of the embryo. Finally, two anchor points are selected along this axis at $y = y_a$ and $y = y_b$. These anchors represent the computed intersections of the lateral 2D symmetry plane with edge of the embryo and are subsequently used to compute the germ band deviation (see "Germ band deviation measurements").

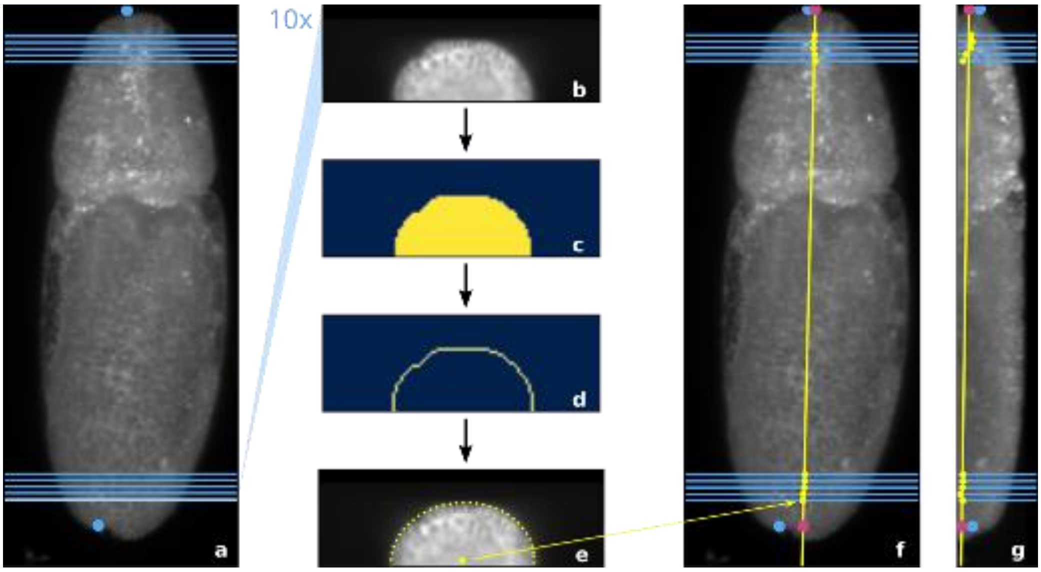


**Suppl. Methods Figure 2:** (a) Z-projection of a 3D image stack of a *Drosophila* embryo. Blue dots indicate the manually identified micropyle and the most posterior germ band point. Blue lines mark ten planes where XZ-slices are extracted from the image. (b) An example XZ-slice from the 3D stack. (c) The binary mask generated using Otsu's thresholding method. (d) Border pixels extracted from the binary mask. (e) An ellipse fitted to the border pixels, with its center represented by a yellow dot. (f) Z-projections of the embryo, yellow dots marking the fitted ellipse centers from ten extracted XY-slices. The anterior-posterior axis is indicated by the yellow line, fitted through the ellipse centers. Pink circles represent the extracted anterior and posterior anchor points. (g) Same as (f) but projected along the X-axis.

Note: This method for identifying the anterior and posterior anchor was chosen for its robustness in handling incomplete image data on the dorsal side. In some 3D image stacks, the anterior tip of the embryo is not visible; however, this approach reliably determines the anterior anchor. The ten XZ-slices are positioned near the embryo's tips to minimize instability in the central region, where substantial tissue movement occurs. However, slices close to the tips are also avoided, as the shallow angle between the slice and the embryo surface otherwise causes instability.
