## Supplemental Note for "“Embryo-eggshell interaction counteracts chiral bias in early *Drosophila* morphogenesis”"

This Supplemental Note divides into three parts: In the first part, we derive the mechanical model, used in the main text, that treats the germband as a pushed elastic line in the plane. In the second part, we analyse an alternative model that treats the germband as a growing, rather than pushed elastic line. In the final part, we discuss an extension that takes the "pushed" model from the plane to the surface of an ellipsoid.

### I. "PUSHED" MODEL OF GERMBAND EXTENSION

As discussed in the main text, we model the germband midline as an effective pushed inextensible elastic line of length $L$ in the plane. The undeformed configuration of the line is straight. Let $\boldsymbol{r}(s,t)$ denote the position of a point of this line, where $s \in [0, L]$ is arclength along the line, and $t$ is time. Using subscripts to denote partial derivatives, the velocity of the line is thus $\boldsymbol{r}_t$. Let $\boldsymbol{t}$ and $\boldsymbol{n}$ be the tangent and normal vectors to the line, which satisfy $\boldsymbol{t} = \boldsymbol{r}_s$ and $\boldsymbol{t}_s = -\kappa\boldsymbol{n}$, where $\kappa$ is the curvature of the line. The external forces acting on the line are

(1) a force at the tip of germband, $s = L$, representing the resistance to germband extension from Scab attachment and the hindgut invagination at the tip of the germband,

$$\boldsymbol{f}_{\text{tip}} = -(\zeta_{\|}\boldsymbol{tt} + \zeta_{\perp}\boldsymbol{nn}) \cdot \boldsymbol{r}_t, \tag{S1}$$

with $\zeta_{\|}$ and $\zeta_{\perp}$ denoting the resistive coefficients in the tangential and normal directions;

(2) forces along the line, representing the resistance to germband extension from the surrounding tissues and the vitelline,

$$\boldsymbol{f}_{\text{friction}} = -(\gamma_{\|}\boldsymbol{tt} + \gamma_{\perp}\boldsymbol{nn}) \cdot \boldsymbol{r}_t, \tag{S2}$$

where $\gamma_{\|}$ and $\gamma_{\perp}$ are the respective resistive coefficients in the tangential and normal directions to the line;

(3) a pushing force at the beginning of the germband, $s = 0$, representing the cell intercalations that drive germband extension,

$$\boldsymbol{f}_0 = F\boldsymbol{t}. \tag{S3}$$

---

[a] Present address: Institut Pasteur, 25-28 Rue du Dr Roux, 75015 Paris, France

[*] These authors contributed equally to this work.

[b] Present address: European Molecular Biology Laboratory Barcelona, Carrer del Doctor Aiguader 88, 08003 Barcelona, Spain

#### A. Derivation of the governing equations

The bending energy of an elastic line is

$$\mathcal{E} = \frac{A}{2}\int_0^L \kappa(s,t)^2\,\mathrm{d}s, \tag{S4}$$

where $A$ is the bending modulus and $\kappa = \|\boldsymbol{r}_{ss}\|$ is the curvature. We impose inextensibility, $\|\boldsymbol{r}_s\| = 1$, by introducing a Lagrange multiplier function $\Lambda(s,t)$ that represents the tension in the germband. The Lagrangian for the problem is thus

$$\mathcal{L} = \int_0^L \left(\frac{A}{2}\|\boldsymbol{r}_{ss}\|^2 - \frac{\Lambda}{2}\|\boldsymbol{r}_s\|^2\right)\mathrm{d}s. \tag{S5}$$

The following derivation is similar to calculations in Ref. [S1]. The variation of this Lagrangian is

$$\begin{aligned}\delta\mathcal{L} = {}&\left[\!\left[ A\boldsymbol{r}_{ss}\cdot\delta\boldsymbol{r}_s - (A\boldsymbol{r}_{sss} + \Lambda\boldsymbol{r}_s)\cdot\delta\boldsymbol{r}\right]\!\right]_0^L \\ &+ \int_0^L \underbrace{\left[A\boldsymbol{r}_{ssss} + (\Lambda\boldsymbol{r}_s)_s\right]}_{-\boldsymbol{f}_{\text{elastic}}}\cdot\delta\boldsymbol{r}\,\mathrm{d}s.\end{aligned} \tag{S6}$$

Meanwhile, the infinitesimal work done by the resistive forces on the germband midline is

$$\delta\mathcal{W} = \int_0^L \boldsymbol{f}_{\text{friction}}\cdot\delta\boldsymbol{r}\,\mathrm{d}s + \boldsymbol{f}_{\text{tip}}\cdot\delta\boldsymbol{r}(L) + \boldsymbol{f}_0\cdot\delta\boldsymbol{r}(0). \tag{S7}$$

Imposing $\delta\mathcal{L} = \delta\mathcal{W}$ leads to

$$A\boldsymbol{r}_{ssss} + (\Lambda\boldsymbol{r}_s)_s + (\gamma_{\|}\boldsymbol{tt} + \gamma_{\perp}\boldsymbol{nn})\cdot\boldsymbol{r}_t = \boldsymbol{0}, \tag{S8}$$

subject to the torque-free boundary conditions [S2]

$$\boldsymbol{r}_{ss}(0,t) = \boldsymbol{r}_{ss}(L,t) = \boldsymbol{0} \tag{S9}$$

and to the force balances at the ends of the germband,

$$A\boldsymbol{r}_{sss}(0,t) + \Lambda(0,t)\boldsymbol{r}_s(0,t) - \boldsymbol{f}_0 = \boldsymbol{0}, \tag{S10a}$$

$$A\boldsymbol{r}_{sss}(L,t) + \Lambda(L,t)\boldsymbol{r}_s(L,t) + \boldsymbol{f}_{\text{tip}} = \boldsymbol{0}. \tag{S10b}$$

To determine the velocity $\boldsymbol{r}_t = (\boldsymbol{r}_t\cdot\boldsymbol{t})\boldsymbol{t} + (\boldsymbol{r}_t\cdot\boldsymbol{n})\boldsymbol{n}$ of the germband midline, we compute, from Eq. (S8),

$$\boldsymbol{r}_t\cdot\boldsymbol{t} = -\frac{1}{\gamma_{\|}}\left[A\boldsymbol{r}_{ssss} + (\Lambda\boldsymbol{r}_s)_s\right]\cdot\boldsymbol{t}, \tag{S11a}$$

$$\boldsymbol{r}_t\cdot\boldsymbol{n} = -\frac{1}{\gamma_{\perp}}\left[A\boldsymbol{r}_{ssss} + (\Lambda\boldsymbol{r}_s)_s\right]\cdot\boldsymbol{n}. \tag{S11b}$$

With $\boldsymbol{r}_s = \boldsymbol{t}$ and $\boldsymbol{t}_s = -\kappa\boldsymbol{n}$, $\boldsymbol{n}_s = \kappa\boldsymbol{t}$, we further compute

$$\boldsymbol{r}_{ss} = -\kappa\boldsymbol{n}, \tag{S12a}$$
$$\boldsymbol{r}_{sss} = -\kappa_s\boldsymbol{n} - \kappa^2\boldsymbol{t}, \tag{S12b}$$
$$\boldsymbol{r}_{ssss} = \left(-\kappa_{ss} + \kappa^3\right)\boldsymbol{n} - 3\kappa\kappa_s\boldsymbol{t}. \tag{S12c}$$

It follows that

$$\boldsymbol{r}_t = \frac{1}{\gamma_\perp}\left[A\left(\kappa_{ss} - \kappa^3\right) + \kappa\Lambda\right]\boldsymbol{n} + \frac{1}{\gamma_\parallel}(3A\kappa\kappa_s - \Lambda_s)\boldsymbol{t}. \tag{S13}$$

Differentiating this equation,

$$\boldsymbol{r}_{st} = \left\{\frac{1}{\gamma_\perp}\left[A\left(\kappa_{sss}-3\kappa^2\kappa_s\right) + \kappa_s\Lambda + \kappa\Lambda_s\right] - \frac{1}{\gamma_\parallel}\left(3A\kappa^2\kappa_s - \kappa\Lambda_s\right)\right\}\boldsymbol{n} + \left\{\frac{1}{\gamma_\perp}\left[A\left(\kappa\kappa_{ss}-\kappa^4\right) + \kappa^2\Lambda\right] + \frac{1}{\gamma_\parallel}\left[3A\left(\kappa_s^2+\kappa\kappa_{ss}\right)-\Lambda_{ss}\right]\right\}\boldsymbol{t}. \tag{S14}$$

Let $\theta(s,t)$ be the tangent angle to the germband midline, so that $\boldsymbol{t} = (\cos\theta, \sin\theta)$ and $\boldsymbol{n} = (\sin\theta, -\cos\theta)$, and hence $\theta_s = \kappa$. This implies that $\boldsymbol{r}_{st} = \boldsymbol{t}_t = -\theta_t\boldsymbol{n}$, whence, by comparison with Eq. (S14),

$$\theta_t = -\frac{1}{\gamma_\perp}\left[A\left(\theta_{ssss} - 3\theta_s^2\theta_{ss}\right) + \theta_{ss}\Lambda + \theta_s\Lambda_s\right] + \frac{1}{\gamma_\parallel}\left(3A\theta_s^2\theta_{ss} - \theta_s\Lambda_s\right), \tag{S15}$$

and

$$0 = \frac{1}{\gamma_\perp}\left[A\theta_s\left(\theta_{sss} - \theta_s^3\right) + \theta_s^2\Lambda\right] + \frac{1}{\gamma_\parallel}\left[3A\left(\theta_{ss}^2 + \theta_s\theta_{sss}\right) - \Lambda_{ss}\right]. \tag{S16}$$

Using Eq. (S12a) and with $\theta_s = \kappa$, the torque-free boundary conditions (S9) become [S2]

$$\theta_s(0,t) = 0, \qquad \theta_s(L,t) = 0. \tag{S17}$$

Moreover, using definition (S3) and Eq. (S12b), Eq. (S10a) yields

$$-A\theta_{ss}(0,t)\boldsymbol{n} - A\theta_s(0,t)^2\boldsymbol{t} + \Lambda(0,t)\boldsymbol{t}(0,t) = F\boldsymbol{t}, \tag{S18}$$

from which, using the first of Eqs. (S17), we obtain two more boundary conditions,

$$\theta_{ss}(0,t) = 0, \qquad \Lambda(0,t) = F. \tag{S19}$$

Similarly, from Eq. (S10b), we find using Eq. (S12b), that

$$-A\theta_s^2(L,t) = -\Lambda(L,t) + \zeta_\parallel(\boldsymbol{t}\cdot\boldsymbol{r}_t), \tag{S20a}$$
$$-A\theta_{ss}(L,t) = \zeta_\perp(\boldsymbol{n}\cdot\boldsymbol{r}_t). \tag{S20b}$$

Thence, using Eq. (S13) and the first of Eqs. (S17) again,

$$0 = -\Lambda(L,t) - \frac{\zeta_\parallel}{\gamma_\parallel}\Lambda_s(L,t), \tag{S21a}$$
$$-\theta_{ss}(L,t) = \frac{\zeta_\perp}{\gamma_\perp}\theta_{sss}(L,t), \tag{S21b}$$

which provide the final two boundary conditions for Eqs. (S15) and (S16).

*Nondimensionalisation*

We nondimensionalise the equations governing the shape of the germband midline and their boundary conditions by setting

$$s = \lambda\hat{s}, \quad L = \lambda\hat{L}, \quad t = \tau\hat{t}, \quad \Lambda = \not{f}\hat{\Lambda}, \quad F = \not{f}\hat{F}, \tag{S22}$$

where the dimensional scalings are

$$\lambda = \frac{\zeta_\perp}{\gamma_\perp}, \qquad \tau = \frac{\zeta_\perp^4}{A\gamma_\perp^3}, \qquad \not{f} = \frac{A\gamma_\perp^2}{\zeta_\perp^2}, \tag{S23}$$

and the hats denote dimensionless variables. We introduce $\ell = \hat{L}$, $f = \hat{F}$ for use in the main text. Dropping the remaining hats for convenience, the dimensionless forms of Eqs. (S15) and (S16) are

$$\theta_t = -\theta_{ssss} - \theta_{ss}\Lambda - \theta_s\left(\Lambda_s - 3\theta_s\theta_{ss}\right)\left(1 + \frac{\gamma_\perp}{\gamma_\parallel}\right), \tag{S24}$$

and

$$\Lambda_{ss} = 3\left(\theta_{ss}^2 + \theta_s\theta_{sss}\right) + \frac{\gamma_\parallel}{\gamma_\perp}\left[\theta_s\left(\theta_{sss} - \theta_s^3\right) + \theta_s^2\Lambda\right]. \tag{S25}$$

They are subject to the dimensionless forms of the boundary conditions (S17), (S19), and (S21), which are

$$\theta_s(0,t) = 0, \qquad \theta_s(\ell,t) = 0, \tag{S26a}$$
$$\theta_{ss}(0,t) = 0, \qquad \theta_{ss}(\ell,t) + \theta_{sss}(\ell,t) = 0, \tag{S26b}$$

and

$$\Lambda(0,t) = f, \qquad \Lambda(\ell,t) + \eta a\Lambda_s(\ell,t) = 0, \tag{S27}$$

where

$$a = \frac{\zeta_\parallel}{\zeta_\perp}, \qquad \eta = \frac{\gamma_\perp}{\gamma_\parallel}. \tag{S28}$$

#### B. Normal-mode analysis of buckling

We analyse the buckling problem by performing a linear stability analysis. On linearising Eq. (S25), we obtain

$$\Lambda_{ss} = 0, \tag{S29}$$

which, with boundary conditions (S27), integrates to

$$\Lambda(s) = f\left(1 - \frac{s}{\eta a + \ell}\right). \tag{S30}$$

On substituting this result into Eq. (S24) and linearising, we then find

$$\theta_t = -\theta_{ssss} - f\left(1 - \frac{s}{\eta a + \ell}\right)\theta_{ss} + \frac{(1+\eta)f}{\eta a + \ell}\theta_s, \tag{S31}$$

To solve this equation subject to its boundary conditions (S26), we make the normal mode ansatz $\theta(s,t) = \Theta(s)\mathrm{e}^{\omega t}$ to separate the time and space dependence of the shape of the germband. This reduces the stability problem to the eigenvalue problem

$$\Theta_{ssss} + f\left(1 - \frac{s}{\eta a + \ell}\right)\Theta_{ss} - \frac{(1+\eta)f}{\eta a + \ell}\Theta_s + \omega\Theta = 0, \tag{S32}$$

subject to

$$\Theta_s(0) = \Theta_s(\ell) = \Theta_{ss}(0) = \Theta_{ss}(\ell) + \Theta_{sss}(\ell) = 0, \tag{S33}$$

from Eqs. (S26). The eigenvalue $\omega$ determines the stability of the germband line: if $\mathrm{Re}(\omega) < 0$, small perturbations decay and the germband remains straight; if $\mathrm{Re}(\omega) > 0$, small perturbations grow and the germband bends. The instability in the latter case grows monotonically if $\mathrm{Im}(\omega) = 0$; it is is an oscillating "flutter" instability if $\mathrm{Im}(\omega) \neq 0$. The stability diagram therefore depends on the dimensionless pushing force $f$, the dimensionless length of the germband $\ell$, and the dimensionless friction ratios $a$ and $\eta$.

While the governing equations (S15) and (S16) are equivalent to equations derived in Ref. [S1], our normal-mode equation (S32) is different, and also different from normal-mode equations found, e.g., in Refs. [S3, S4], because the boundary conditions of our problem are different.

***Numerical solution of the normal-mode equation***

Numerical solution of Eq. (S32) is not straightforward because this normal-mode equation is not autonomous. We determine the eigenvalue $\omega$ for general values of the parameters $f, \ell, a, \eta$ in the following way:

(1) We first solve the equations semi-analytically for $f \ll 1$, in which limit Eq. (S32) becomes

$$\Theta_{ssss} + \omega\Theta = 0, \tag{S34}$$

the general solution of which is

$$\Theta(s) = c_1 \cos \Omega s + c_2 \sin \Omega s + c_3 \cosh \Omega s + c_4 \sinh \Omega s, \tag{S35}$$

with $\Omega = (-\omega)^{1/4}$, and where $c_1, c_2, c_3, c_4$ are constants of integration. Their values are determined by substituting this solution in the boundary conditions (S33). These conditions can be written as

$$\boldsymbol{M}(\Omega, \ell)\boldsymbol{c} = \boldsymbol{0}, \tag{S36}$$

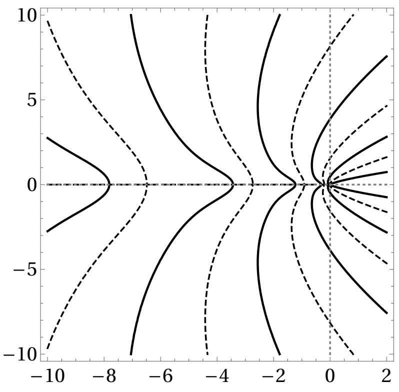


FIG. S1. Contour plot of $\mathrm{Re}(m) = 1$ (solid lines) and $\mathrm{Im}(m) = 0$ (dashed lines) in the $(\mathrm{Re}(\omega), \mathrm{Im}(\omega))$ plane for $\ell = 10$, showing that all solutions of $m(\Omega; \ell) = 0$ have $\omega = -\Omega^4 \leqslant 0$. Dotted lines show $\mathrm{Re}(\omega) = 0$ and $\mathrm{Im}(\omega) = 0$.

where $\boldsymbol{c} = (c_1, c_2, c_3, c_4)$, and $\boldsymbol{M}$ is a matrix with entries depending on $\Omega$ and $\ell$. These linear equations have a non-trivial solution if and only if $\det \boldsymbol{M}(\Omega, \ell) = 0$. Using Mathematica (Wolfram, Inc.), we find that this condition reduces to the transcendental equation

$$m(\Omega; \ell) \equiv \Omega(\cos \Omega\ell \sinh \Omega\ell - \cosh \Omega\ell \sin \Omega\ell) + \cos \Omega\ell \cosh \Omega\ell = 1. \tag{S37}$$

Numerical contour plots (Fig. S1) of $\mathrm{Re}(m) = 1$ and $\mathrm{Im}(m) = 0$ in the $(\mathrm{Re}(\omega), \mathrm{Im}(\omega))$ plane suggest that all solutions have $\omega \leqslant 0$, i.e. $\Omega \geqslant 0$. These solutions can be found using the `fsolve` function of Matlab (The MathWorks, Inc.), starting from initial guesses obtained from approximate solutions for $|\Omega| \gg 1$. In this limit,

$$1 = m(\Omega; \ell) \approx \frac{\mathrm{e}^{\Omega\ell}}{2}\left[\Omega(\cos \Omega\ell - \sin \Omega\ell) + \cos \Omega\ell\right] + \mathcal{O}\left(\mathrm{e}^{-\Omega\ell}\right), \tag{S38}$$

which can only hold if $\cos \Omega\ell + \Omega(\cos \Omega\ell - \sin \Omega\ell) \approx 0$ which means $\tan \Omega\ell \approx 1 + 1/\Omega \approx 1$ i.e., only if $\Omega\ell \approx (n + 1/4)\pi$ for $n = 0, 1, 2, \ldots$. However, the exact solution corresponding to the approximate solution with $n = 0$ turns out to be $\Omega = 0$, which we reject. Therefore $\omega = -\Omega^4 \approx -[(n + 1/4)\pi/\ell]^4$ for $n = 1, 2, \ldots$.

(2) We then solve the full normal-mode equation (S32) by using the principle of superposition to write the solution that satisfies boundary conditions (S33) in the form

$$\Theta(s) = c_1\Theta^{(1)}(s) + c_2\Theta^{(2)}(s) + c_3\Theta^{(3)}(s) + c_4\Theta^{(4)}(s), \tag{S39}$$

where $c_1, c_2, c_3, c_4$ are yet again constants of integration, $\Theta^{(1)}, \Theta^{(2)}, \Theta^{(3)}, \Theta^{(4)}$ are linearly independent solutions satisfying the boundary conditions

$$\Theta^{(1)}(0) = 1,\ \Theta_s^{(1)}(0) = 0,\ \Theta_{ss}^{(1)}(0) = 0,\ \Theta_{sss}^{(1)}(0) = 0, \tag{S40a}$$

$$\Theta^{(2)}(0) = 0,\ \Theta_s^{(2)}(0) = 1,\ \Theta_{ss}^{(2)}(0) = 0,\ \Theta_{sss}^{(2)}(0) = 0, \tag{S40b}$$

$$\Theta^{(3)}(0) = 0,\ \Theta_s^{(3)}(0) = 0,\ \Theta_{ss}^{(3)}(0) = 1,\ \Theta_{sss}^{(3)}(0) = 0, \tag{S40c}$$

$$\Theta^{(4)}(0) = 0,\ \Theta_s^{(4)}(0) = 0,\ \Theta_{ss}^{(4)}(0) = 0,\ \Theta_{sss}^{(4)}(0) = 1. \tag{S40d}$$

By construction, the Wronskian of these solutions is $W = 1$; since it is nonzero, the solutions are indeed linearly independent. We generate these linearly independent solutions numerically using the `ode45` solver of Matlab. Imposing the boundary conditions again leads to a determinant condition $\det \boldsymbol{M}(\omega, f, \ell, a, \eta) = 0$. Numerically, we remove the trivial solution $\omega = 0$ by solving $\omega^{-1} \det \boldsymbol{M}(\omega, f, \ell, a, \eta) = 0$ using the `fsolve` function of Matlab, for increasing values of $f$, by starting from small values and using the solutions obtained in (1) above as initial guesses.

In this way, for each set of parameter values, we can find all eigenvalues by starting from the complete spectrum generated in (1) above; numerically, we track multiple eigenvalues obtained in this way to find the eigenvalue with maximal real part, and hence determine the onset of instability.

#### *Effect of $\eta$ on the stability diagram*

In the main text, we show results for $\eta = 2$, which is the bulk friction anisotropy ratio of resistive force theory [S5], but our model is of course only one of an effective elastic line. Still, other values of $\eta$ do not change the stability diagram qualitatively (Fig. S2). At a quantitative level, the region of parameter space in which the germband is unstable is slightly larger for smaller values of $\eta$ (Fig. S2A) compared to larger values (Fig. S2B).

#### *Discontinuity in the stability diagram*

The stability diagrams in the main text and in Fig. S2 display a discontinuity at the transition between non-flutter and flutter instability. This is *not* a numerical artefact, but a true discontinuity. To explain this discontinuity, we consider plots of $\mathrm{Re}(\omega), \mathrm{Im}(\omega)$ against $f$, at fixed $\ell, a, \eta$. Let $f_{\mathrm{Re}}$ denote the value of $f$ at which $\mathrm{Re}(\omega)$ first becomes positive, and let $f_{\mathrm{Im}}$ denote the value of $f$ at which $\mathrm{Im}(\omega)$ first becomes nonzero. Two qualitatively different diagrams can arise, depending on whether $f_{\mathrm{Re}} > f_{\mathrm{Im}}$ (Fig. S3A) or $f_{\mathrm{Re}} < f_{\mathrm{Im}}$ (Fig. S3B).

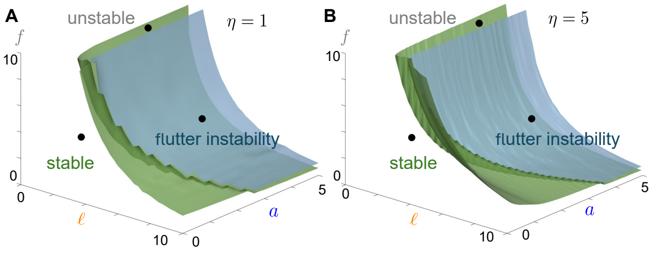


FIG. S2. Effect of $\eta$ on the stability diagram. Phase diagrams of the instability in $(\ell, a, f)$ space, showing the regions of parameter space in which the germband is stable, unstable, and shows a flutter instability respectively, for (A) $\eta = 1$ and (B) $\eta = 5$. The two phase diagrams show that $\eta$ affects the instability quantitatively, but not qualitatively.

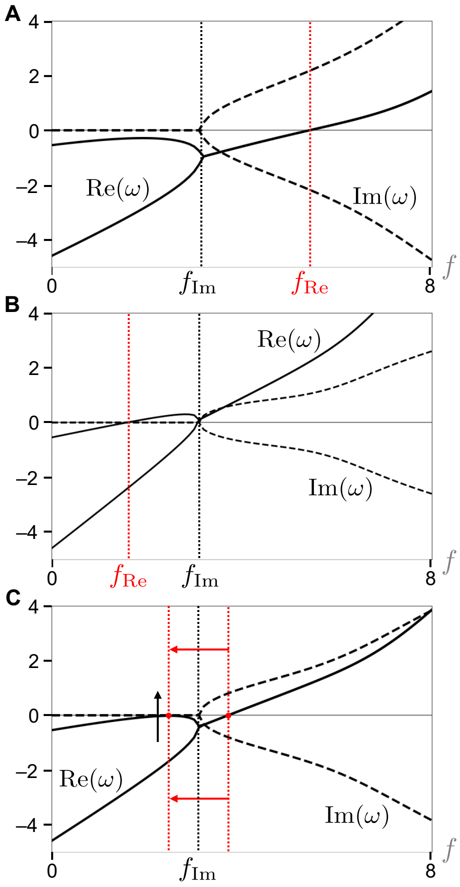


FIG. S3. Discontinuity in the stability diagram. Plots of $\mathrm{Re}(\omega)$ and $\mathrm{Im}(\omega)$ against $f$, at fixed $\ell, a, \eta$ are qualitatively different in the cases (A) $f_{\mathrm{Re}} > f_{\mathrm{Im}}$ and (B) $f_{\mathrm{Re}} < f_{\mathrm{Im}}$, where $f_{\mathrm{Re}}$ is the value of $f$ at which $\mathrm{Re}(\omega)$ first becomes positive and $f_{\mathrm{Im}}$ is the value of $f$ at which $\mathrm{Im}(\omega)$ first becomes non-zero. (C) At the transition between these two cases, i.e., at the transition between flutter and non-flutter instability, $f_{\mathrm{Re}}$ jumps discontinuously. Parameter values: $\ell = 5$, $\eta = 2$, and (A) $a = 0.20$, (B) $a = 7.64$, (C) $a = 2.61$.

The transition between these cases corresponds to the transition between non-flutter and flutter instability. At this transition (Fig. S3C), $f_{\mathrm{Re}}$ jumps discontinuously, as another root of $\mathrm{Re}(\omega) = 0$ appears at the local maximum of $\mathrm{Re}(\omega)$ as a function of $f$. This explains the observed discontinuity.

### II. "GROWING" MODEL OF GERMBAND EXTENSION

An alternative model of germband extension represents the germband midline as a growing, inextensible elastic line, similarly to a model developed in Ref. [S6] for filament extrusion. In other words, the cell intercalations driving germband extension now appear as growth rather than a pushing force. We denote by $U$ the constant velocity at which the germband is growing, so that its length is $L(t) = L_0 + Ut$, where $t$ is time and $L_0$ is the initial length of the germband.

In this model (Fig. S4), the external forces exerting on the germband are, again,

(1) a force at the tip of germband, $s = L(t)$, that represents the resistance to germband extension from Scab attachment

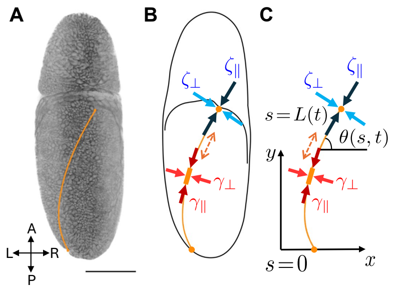


FIG. S4. “Growing” model of germband extension. (A) Dorsal view of a *Drosophila* embryo during germband extension. Orange line: germband midline. Embryo axes: A(nterior), P(osterior), L(eft), R(ight). Scale bar: 100 µm. (B) Mechanical model of the germband midline as a growing inextensible elastic line, with growth representing germband extension. Bulk forces along the germband midline with perpendicular and parallel friction coefficients $\gamma_\perp$, $\gamma_\parallel$ represent friction forces from the surrounding tissues and the vitelline envelope. Tip forces with perpendicular and parallel friction coefficients $\zeta_\perp$, $\zeta_\parallel$ represent friction from the tip attachment of the germband. (C) Two-dimensional projection of the model into the $(x, y)$ plane. Arclength along the inextensible germband midline is $s$, with $s = 0$ corresponding to the posterior end and $s = L(t)$ to the tip of the germband. The tangent angle to the germband is $\theta(s, t)$.

and the hindgut invagination at the tip of the germband,

$$\boldsymbol{f}_{\text{tip}} = -(\zeta_\parallel \boldsymbol{tt} + \zeta_\perp \boldsymbol{nn}) \cdot \frac{\mathrm{D}\boldsymbol{r}}{\mathrm{D}t}, \tag{S41}$$

with $\zeta_\parallel$ and $\zeta_\perp$ denoting, again, the resistive coefficients in the tangential and normal directions;

(2) the friction forces, all along the germband midline, from the surrounding tissue,

$$\boldsymbol{f}_{\text{friction}} = -(\gamma_\parallel \boldsymbol{tt} + \gamma_\perp \boldsymbol{nn}) \cdot \frac{\mathrm{D}\boldsymbol{r}}{\mathrm{D}t}, \tag{S42}$$

The resistive coefficients in tangential and normal directions are $\gamma_\parallel$ and $\gamma_\perp$ respectively.

Here $\boldsymbol{t}$ and $\boldsymbol{n}$ are, again, the tangent and normal vectors to the germband midline. By contrast to the previous “pushed” model, the velocity that appears in these forces is the complete time derivative of position $\boldsymbol{r}$ in the lab frame,

$$\frac{\mathrm{D}\boldsymbol{r}}{\mathrm{D}t} = \frac{\partial \boldsymbol{r}}{\partial t} + \frac{\partial s}{\partial t}\boldsymbol{t}. \tag{S43a}$$

In this expression, the grown arclength $s(t)$ is related to the initial arclength $s_0$ by $s(t)/s_0 = L(t)/L_0 = 1 + Ut/L_0$, assuming uniform growth. Hence $\partial s/\partial t = Us_0/L_0 = Us/(L_0 + Ut)$. Thus

$$\frac{\mathrm{D}\boldsymbol{r}}{\mathrm{D}t} = \frac{\partial \boldsymbol{r}}{\partial t} + \frac{Us}{L_0 + Ut}\boldsymbol{t}. \tag{S43b}$$

#### A. Derivation of the governing equations

The elastic force is unchanged from the previous model, and is still given by

$$\boldsymbol{f}_{\text{elastic}} = -A\boldsymbol{r}_{ssss} - (\Lambda \boldsymbol{r}_s)_s. \tag{S44}$$

The force balance $\boldsymbol{f}_{\text{elastic}} + \boldsymbol{f}_{\text{friction}} = \boldsymbol{0}$ now leads to

$$\frac{\mathrm{D}\boldsymbol{r}}{\mathrm{D}t} = \frac{1}{\gamma_\perp}\left[A\left(\kappa_{ss} - \kappa^3\right) + \kappa\Lambda\right]\boldsymbol{n} + \frac{1}{\gamma_\parallel}(3A\kappa_s\kappa - \Lambda_s)\boldsymbol{t}. \tag{S45}$$

As in the “pushed” model, from the $\delta\mathcal{L} = \delta\mathcal{W}$ of the variation of the Lagrangian and the work done by the external force, we obtain the torque-free boundary conditions

$$\boldsymbol{r}_s(0, t) \times \boldsymbol{r}_{ss}(0, t) = \boldsymbol{r}_s(L(t), t) \times \boldsymbol{r}_{ss}(L(t), t) = \boldsymbol{0}, \tag{S46}$$

and the force balances at the end of the germband

$$A\boldsymbol{r}_{sss}(L(t), t) + \Lambda(L(t), t)\boldsymbol{r}_s(L(t), t) + \boldsymbol{f}_{\text{tip}} = \boldsymbol{0}, \tag{S47a}$$

$$A\boldsymbol{r}_{sss}(0, t) + \Lambda(0, t)\boldsymbol{r}_s(0, t) = \boldsymbol{0}. \tag{S47b}$$

##### *Nondimensionalisation*

It is convenient to nondimensionalise these governing equations and their boundary conditions using a different nondimensionalisation from that used for the “pushed” model. We choose to write

$$s = \lambda\hat{s}, \quad L = \lambda\hat{L}, \quad L_0 = \lambda\hat{L}_0, \quad t = \tau\hat{t}, \quad \Lambda = f\hat{\Lambda}, \tag{S48}$$

where the dimensional scalings are

$$\lambda = \left(\frac{A}{\gamma_\parallel U}\right)^{1/3}, \qquad \tau = \frac{\lambda}{U}, \qquad f = \frac{A}{\lambda^2}, \tag{S49}$$

and the hats denote dimensionless variables. Again, we drop hats but use $\ell = \hat{L}$, $\ell_0 = \hat{L}_0$ for consistency with the main text. The dimensionless form of the governing equation (S45) is therefore

$$\frac{\mathrm{D}\boldsymbol{r}}{\mathrm{D}t} = \eta^{-1}\left(\kappa_{ss} - \kappa^3 + \kappa\Lambda\right)\boldsymbol{n} + (3\kappa_s\kappa - \Lambda_s)\boldsymbol{t}, \tag{S50}$$

and boundary conditions (S46) and (S47) become

$$\boldsymbol{r}_s(0, t) \times \boldsymbol{r}_{ss}(0, t) = \boldsymbol{r}_s(\ell(t), t) \times \boldsymbol{r}_{ss}(\ell(t), t) = \boldsymbol{0}, \tag{S51a}$$

$$\boldsymbol{r}_{sss}(\ell(t), t) + \Lambda(\ell(t), t)\boldsymbol{r}_s(\ell(t), t) = \Gamma\left(\boldsymbol{tt} + a^{-1}\boldsymbol{nn}\right) \cdot \frac{\mathrm{D}\boldsymbol{r}}{\mathrm{D}t}, \tag{S51b}$$

$$\boldsymbol{r}_{sss}(0, t) + \Lambda(0, t)\boldsymbol{r}_s(0, t) = \boldsymbol{0}. \tag{S51c}$$

where

$$\eta = \frac{\gamma_\perp}{\gamma_\parallel}, \qquad a = \frac{\zeta_\parallel}{\zeta_\perp}, \qquad \Gamma = \frac{\zeta_\parallel}{\gamma_\parallel \lambda} = \left(\frac{\zeta_\parallel^3 U}{\gamma_\parallel^2 A}\right)^{1/3}. \tag{S52}$$

##### *Derivation of equations for $\Lambda(s, t)$ and $\theta(s, t)$*

In dimensionless form $\partial s/\partial t = s/(\ell_0 + t)$. The dimensionless form of the total derivative is therefore

$$\frac{\mathrm{D}}{\mathrm{D}t} = \frac{\partial}{\partial t} + \frac{s}{\ell_0 + t}\frac{\partial}{\partial s}. \tag{S53}$$

Hence

$$\frac{\mathrm{D}\boldsymbol{r}_s}{\mathrm{D}t} = \boldsymbol{r}_{st} + \frac{s}{\ell_0 + t}\boldsymbol{r}_{ss}, \tag{S54a}$$

$$\frac{\partial}{\partial s}\left(\frac{\mathrm{D}\boldsymbol{r}}{\mathrm{D}t}\right) = \frac{\partial}{\partial s}\left(\boldsymbol{r}_t + \frac{s}{\ell_0 + t}\boldsymbol{r}_s\right) = \boldsymbol{r}_{st} + \frac{s}{\ell_0 + t}\boldsymbol{r}_{ss} + \frac{\boldsymbol{r}_s}{\ell_0 + t}. \tag{S54b}$$

These imply that

$$\frac{\mathrm{D}\boldsymbol{t}}{\mathrm{D}t}=\frac{\partial}{\partial s}\left(\frac{\mathrm{D}\boldsymbol{r}}{\mathrm{D}t}\right)-\frac{\boldsymbol{t}}{\ell_0+t}. \tag{S55}$$

Now, from Eq. (S50), and using $\boldsymbol{t}_s=-\kappa\boldsymbol{n}$, $\boldsymbol{n}_s=\kappa\boldsymbol{t}$ again, we compute

$$\frac{\partial}{\partial s}\left(\frac{\mathrm{D}\boldsymbol{r}}{\mathrm{D}t}\right)=\left\{\eta^{-1}\left[\kappa_{sss}-3\kappa^2\kappa_s+(\kappa\Lambda)_s\right]-\kappa(3\kappa\kappa_s-\Lambda_s)\right\}\boldsymbol{n}+\left[\kappa\eta^{-1}\left(\kappa_{ss}-\kappa^3+\kappa\Lambda\right)+3\kappa_s^2+3\kappa\kappa_{ss}-\Lambda_{ss}\right]\boldsymbol{t}. \tag{S56}$$

Again, we introduce the tangent angle $\theta(s,t)$ to the germband midline, so that $\boldsymbol{t}=(\cos\theta,\sin\theta)$ and $\boldsymbol{n}=(\sin\theta,-\cos\theta)$, and hence $\theta_s=\kappa$. Using the total time derivative given in Eq. (S53),

$$\frac{\mathrm{D}\boldsymbol{t}}{\mathrm{D}t}=\frac{\mathrm{D}}{\mathrm{D}t}(\cos\theta,\sin\theta)=\left(\theta_t+\frac{s\theta_s}{\ell_0+t}\theta_s\right)(-\sin\theta,\cos\theta)=-\left(\theta_t+\frac{\kappa s}{\ell_0+t}\right)\boldsymbol{n}. \tag{S57}$$

Inserting this result into Eq. (S55) yields, from the normal and tangential components respectively,

$$-\theta_t-\frac{\kappa s}{\ell_0+t}=\eta^{-1}\left[\kappa_{sss}-3\kappa^2\kappa_s+(\kappa\Lambda)_s\right]-\kappa(3\kappa\kappa_s-\Lambda_s), \tag{S58a}$$

$$\frac{1}{\ell_0+t}=\kappa\eta^{-1}\left(\kappa_{ss}-\kappa^3+\kappa\Lambda\right)+3\kappa_s^2+3\kappa\kappa_{ss}-\Lambda_{ss}. \tag{S58b}$$

Using $\kappa=\theta_s$, this gives the partial differential equation for $\theta(s,t)$,

$$\eta\theta_t=-\theta_{ssss}+\theta_{ss}\left[3(1+\eta)\theta_s^2-\Lambda\right]-(1+\eta)\theta_s\Lambda_s-\frac{\eta s\theta_s}{\ell_0+t}, \tag{S59a}$$

as well as the ordinary differential equation for the $\Lambda(s,t)$,

$$\eta\Lambda_{ss}=\theta_s^2\Lambda-\theta_s^4+(3\eta+1)\theta_s\theta_{sss}+3\eta\theta_{ss}^2-\frac{\eta}{\ell_0+t}. \tag{S59b}$$

#### B. Normal-mode analysis of buckling

To analyse the buckling of the germband midline in this model, we linearise Eqs. (S59b) and (S59a) for $\theta\ll 1$ to obtain

$$\Lambda_{ss}=-\frac{1}{\ell_0+t}, \tag{S60a}$$

$$\eta\theta_t=-\theta_{ssss}-\theta_{ss}\Lambda-\theta_s(1+\eta)\Lambda_s-\frac{\eta s\theta_s}{\ell_0+t}. \tag{S60b}$$

From Eqs. (S51) and using Eqs. (S12a) and (S12b), the linearised dimensionless boundary conditions are

$$\theta_s(0,t)=0,\quad \theta_s(\ell_0+t,t)=0, \tag{S61a}$$

$$\theta_{ss}(0,t)=0,\quad \theta_{ss}(\ell_0+t,t)=-\Gamma(a\eta)^{-1}\theta_{sss}(\ell_0+t,t), \tag{S61b}$$

$$\Lambda(0,t)=0,\quad \Lambda(\ell_0+t,t)=-\Gamma\Lambda_s(\ell_0+t,t), \tag{S61c}$$

where we have used the first two conditions to simplify the final four conditions. We can now solve Eq. (S60a) subject to boundary conditions (S61c) to get

$$\Lambda(s,\tau)=-\frac{s^2}{2(\ell_0+t)}+\frac{\Gamma+\frac{1}{2}(\ell_0+t)}{\Gamma+(\ell_0+t)}s. \tag{S62}$$

Next, to obtain equations that can be solved by a normal mode analysis, we assume that the germband midline is growing slowly to approximate $\ell_0+t\approx\ell_0$ and hence eliminate the explicit dependence on time from these equations. Making the normal mode ansatz $\theta(s,t)=\Theta(s)\mathrm{e}^{\omega t}$ again, Eq. (S60b) becomes

$$\Theta_{ssss}-\Theta_{ss}\left(\frac{s^2}{2\ell_0}-Cs\right)-\Theta_s\left[\frac{s}{\ell_0}-(1+\eta)C\right]+\eta\omega\Theta=0, \tag{S63}$$

where we have introduced

$$C=\frac{\Gamma+\frac{1}{2}\ell_0}{\Gamma+\ell_0}. \tag{S64}$$

Boundary conditions (S61a) and (S61b) become

$$\Theta_s(0)=0,\qquad \Theta_s(\ell_0)=0, \tag{S65a}$$

$$\Theta_{ss}(0)=0,\qquad \Theta_{ss}(\ell_0)=-\beta\Theta_{sss}(\ell_0), \tag{S65b}$$

where $\beta=\Gamma(a\eta)^{-1}$. We stress that, in spite of the "slow growth" assumption, Eq. (S63) has a different form from the normal-mode equation (S32) that we found for the "pushed" model of germband extension.

***Numerical solution of the normal-mode equation***

The final normal-mode equation (S63) and its boundary conditions (S65) thus depend on the dimensionless parameters $\ell_0, C, \beta, \eta$. Again, we solve this using a "Wronskian method":

(1) We first solve the equations semi-analytically for $\beta\ll 1$, $\ell_0\ll 1$, $C\ll 1$. Because $s\leqslant\ell_0$, Eq. (S63) becomes, in this limit,

$$\Theta_{ssss}+\eta\omega\Theta=0. \tag{S66a}$$

We make the change of variable $s=\ell_0 S$, so that

$$\Theta_{SSSS}-\Omega^4\Theta=0, \tag{S66b}$$

where $\Omega^4=-\omega\eta\ell_0^4$. The boundary conditions are

$$\Theta_S(0)=0,\qquad \Theta_S(1)=0, \tag{S67a}$$

$$\Theta_{SS}(0)=0,\qquad \Theta_{SS}(1)=0. \tag{S67b}$$

The solution of Eq. (S66b) is

$$\Theta(S)=c_1\sin\Omega S+c_2\cos\Omega S+c_3\sinh\Omega S+c_4\cosh\Omega S, \tag{S68}$$

where $c_1, c_2, c_3, c_4$ are constants of integration. As in "pushed" model, their values are determined by substituting this solution in the boundary conditions (S67). These boundary conditions can again be written as

$$\boldsymbol{M}(\Omega)\boldsymbol{c}=\boldsymbol{0}, \tag{S69}$$

where $\boldsymbol{c} = (c_1, c_2, c_3, c_4)$, and $\boldsymbol{M}$ is a matrix with entries depending on $\Omega$. These linear equations have a non-trivial solution if and only if $\det \boldsymbol{M}(\Omega) = 0$. Using Mathematica, we find the transcendental equation

$$\cos\Omega \cosh\Omega = 1. \tag{S70}$$

This has a trivial solution $\Omega = 0$. Other solutions have $\Omega \approx (n + 1/2)\pi$, i.e., $\omega \approx -(n+1/2)^4\pi^4/\eta\ell_0^4$, for $n = 1, 2, 3, \ldots$. We determine better approximations of these eigenvalues by solving Eq. (S70) numerically using the `fsolve` function of Matlab.

(2) The next step is to increase $\beta$. The approximate eigenvalue problem is still Eq. (S66b), but the final boundary condition in Eq. (S67b) is now

$$\Theta_{SS}(1) + B\Theta_{SSS}(1) = 0, \tag{S71}$$

where $B = \beta/\ell_0$. This means that condition (S70) is changed to

$$\cos\Omega\cosh\Omega = 1 + B\Omega\bigl(\cosh\Omega\sin\Omega - \cos\Omega\sinh\Omega\bigr). \tag{S72}$$

Its solutions can be found numerically using the `fsolve` function of Matlab, using those found in the previous case as initial guesses and by increasing $\beta$.

(3) The final step is to increase $\ell_0$ for each value of $C$ and $\eta$ by solving the full eigenvalue problem. With the rescalings introduced above Eq. (S63) becomes

$$\Theta_{SSSS} = \ell_0^3\left\{\Theta_{SS}\left(\frac{S^2}{2} - CS\right) + \Theta_S[S - (1+\eta)C]\right\} - \Omega^4\Theta, \tag{S73}$$

and is to be solved subject to

$$\Theta_S(0) = 0, \qquad \Theta_S(1) = 0, \tag{S74a}$$
$$\Theta_{SS}(0) = 0, \qquad \Theta_{SS}(1) = -B\Theta_{SSS}(1). \tag{S74b}$$

We can now solve this problem using the "Wronskian method" introduced for the "pushed" model, using the eigenvalues found above as initial guesses for the numerical solution and increasing $\ell_0$.

##### *Results: Absence of non-flutter instability*

We computed eigenvalues $\omega$ with this numerical method for different values of the dimensionless parameters $\ell_0, B, C, \eta$. As expected, a short enough germband is stable while a longer germband is unstable, as illustrated in Fig. S5. In these computations, we could not however find any parameter values with $\mathrm{Re}(\omega) > 0$ and $\mathrm{Im}(\omega) = 0$, i.e., for which the germband displays a non-flutter instability. While this does not of course provide a mathematical proof that no such parameter values exist, it suggests that the "growing" model of germband extension is less appropriate than the "pushed" model that we analyse in the main text; this is also in agreement, as discussed there, with what is known about the biology of germband extension.

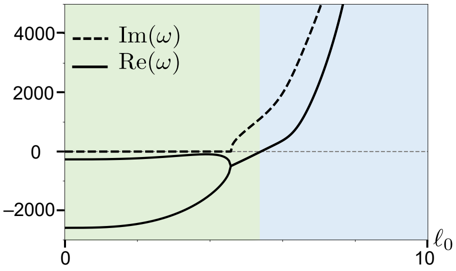


FIG. S5. Absence of non-flutter instability in the "growing" model of germband extension, illustrated by plots of $\mathrm{Re}(\omega)$ and $\mathrm{Im}(\omega)$ against $\ell_0$ for the two leading eigenvalues for $B = 1.005$, $C = 1$, $\eta = 2$. A long enough growing germband is unstable to a flutter instability (blue shaded area), while a short germband is stable (green shaded area).

### III. "PUSHED" MODEL OF GERMBAND EXTENSION ON AN ELLIPSOID

Finally, we discuss how to extend the "pushed" model of germband extension to the surface of an ellipsoid, to mimic the three-dimensional shape of the *Drosophila* embryo and hence describe the effect of embryonic curvature on the buckling instability of the germband. The position of a point on the germband midline, confined to the surface of an ellipsoid with semi-major axis $p$ and semi-minor axis $q \leqslant p$ (Fig. S6) is

$$\boldsymbol{r}(s,t) = \bigl(p\cos\theta(s,t)\cos\phi(s,t), q\sin\theta(s,t), q\cos\theta(s,t)\sin\phi(s,t)\bigr), \tag{S75a}$$

in which $\theta(s,t)$ is the latitude, with $0 \leqslant \theta < 2\pi$, and $\phi(s,t)$ is the azimuthal angle, with $0 \leqslant \phi < 2\pi$, and where $s$ is

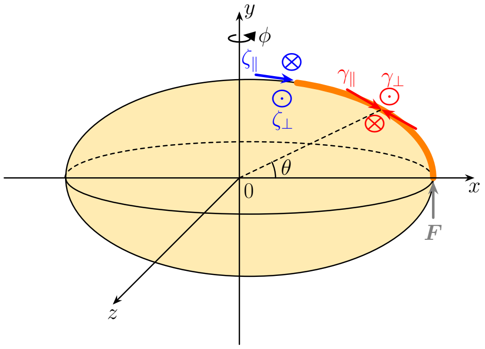


FIG. S6. Mechanical model of the germband midline as an inextensible elastic line on an ellipsoid of semi-major axis $p$ and semi-minor axis $q$. The elastic line is pushed by a force $\boldsymbol{F}$ driving germband extension. Bulk forces along the germband midline with perpendicular and parallel friction coefficients $\gamma_\perp$, $\gamma_\parallel$ represent friction forces from the surrounding tissues and the vitelline envelope. Tip forces with perpendicular and parallel friction coefficients $\zeta_\perp$, $\zeta_\parallel$ represent friction from the tip attachment of the germband. A point on the germband midline in Cartesian axes $(x, y, z)$ is determined by its latitude $\theta(s,t)$ and azimuth $\phi(s,t)$, where $s$ is arclength. The posterior end of the germband $s = 0$ has $\theta = 0$ at $t = 0$. The straight configuration of the germband midline has $\phi = 0$.

arclength. We assume that the initial latitude of the pushed end of the germband is $\theta_0(0,0) = 0$. If the germband is straight, $\theta(s,t) = \theta_0(s,t)$ and $\phi = 0$, corresponding to

$$\boldsymbol{r}(s,t) = \bigl(p\cos\theta_0(s,t), q\sin\theta_0(s,t), 0\bigr). \tag{S75b}$$

Hence

$$s = \int_{\theta_0(0,t)}^{\theta_0(s,t)} \bigl(p^2\sin^2\vartheta + q^2\cos^2\vartheta\bigr)^{1/2}\,\mathrm{d}\vartheta. \tag{S76a}$$

As the germband extends (while remaining straight) and because it is inextensible, each point on the germband moves with constant velocity $u$, to be determined. Thus

$$ut = \int_{\theta_0(s,0)}^{\theta_0(s,t)} \bigl(p^2\sin^2\vartheta + q^2\cos^2\vartheta\bigr)^{1/2}\,\mathrm{d}\vartheta. \tag{S76b}$$

Taking Eq. (S76a) for $t = 0$ and adding it to Eq. (S76b) implies

$$s + ut = \int_{0}^{\theta_0(s,t)} \bigl(p^2\sin^2\vartheta + q^2\cos^2\vartheta\bigr)^{1/2}\,\mathrm{d}\vartheta, \tag{S76c}$$

on recalling $\theta_0(0,0) = 0$. This result can be simplified in terms of the incomplete elliptic integral of the second kind [S7],

$$\mathrm{E}(\theta \mid k) = \int_0^{\theta} \sqrt{1 - k^2\sin^2\vartheta}\,\mathrm{d}\vartheta, \tag{S77}$$

and the corresponding complete integral $\mathrm{E}(k) = \mathrm{E}\bigl(\frac{\pi}{2} \mid k\bigr)$. With this, Eq. (S76c) becomes

$$\frac{s+ut}{p} = \mathrm{E}(k) - \mathrm{E}\bigl(\tfrac{\pi}{2} - \theta_0(s,t) \mid k\bigr) \quad \text{where } k = \sqrt{1 - \frac{q^2}{p^2}}. \tag{S78}$$

Equivalently,

$$\theta_0(s,t) = \frac{\pi}{2} - \mathrm{E}^{-1}\left(\mathrm{E}(k) - \frac{s+ut}{p} \mid k\right). \tag{S79a}$$

This expression is not conducive to further analytical calculations. In what follows we therefore specialise to the spherical case, $q = p$, in which case we have, simply,

$$\theta_0(s,t) = \frac{s+ut}{p}. \tag{S79b}$$

To describe the forces acting on the germband, we introduce a Darboux frame [S8] for the spherical surface in which $\boldsymbol{t}$ is the tangent vector to the midline, $\boldsymbol{n}$ is the normal to the surface, and $\boldsymbol{m} = \boldsymbol{t}\times\boldsymbol{n}$ is the tangent normal. With these definitions, the tip friction, bulk friction, and pushing force on the germband midline are

$$\boldsymbol{f}_{\text{tip}} = -(\zeta_\parallel \boldsymbol{tt} + \zeta_\perp \boldsymbol{mm})\cdot\boldsymbol{r}_t, \tag{S80a}$$
$$\boldsymbol{f}_{\text{friction}} = -(\gamma_\parallel \boldsymbol{tt} + \gamma_\perp \boldsymbol{mm})\cdot\boldsymbol{r}_t, \tag{S80b}$$
$$\boldsymbol{F} = F\boldsymbol{t}, \tag{S80c}$$

by analogy with the forces (S1)–(S3) in the "pushed" model in the plane. In addition to these forces, a reaction force $\boldsymbol{R} \parallel \boldsymbol{n}$ acts along the germband midline to balance the component of the elastic force perpendicular to the surface, so that the germband midline remains confined to the surface.

#### A. Non-dimensional governing equations

Again, we denote by $L$ the length of the germband midline, and impose its inextensibility using a Lagrange multiplier function $\Lambda(s,t)$. The vectorial form of the variation of the Lagrangian remains unchanged from Eq. (S6). We nondimensionalise the problem similarly to the flat case, using again the dimensional scalings (S23). To the nondimensionalisations defined by Eqs. (S22), we must now add

$$p = \lambda\hat{p}, \qquad u = \upsilon\hat{u}, \quad \text{where } \upsilon = \frac{\lambda}{\tau} = \frac{A\gamma_\perp^2}{\zeta_\perp^3}, \tag{S81}$$

using Eqs. (S23). Using $\ell = \hat{L}, f = \hat{F}$ and again dropping hats, and imposing force balance within the surface, we obtain, similarly to the derivation of Eq. (S8),

$$\boldsymbol{r}_{ssss}\cdot\boldsymbol{t} + (\Lambda\boldsymbol{r}_s)_s\cdot\boldsymbol{t} + \eta^{-1}\boldsymbol{r}_t\cdot\boldsymbol{t} = 0, \tag{S82a}$$
$$\boldsymbol{r}_{ssss}\cdot\boldsymbol{m} + (\Lambda\boldsymbol{r}_s)_s\cdot\boldsymbol{m} + \boldsymbol{r}_t\cdot\boldsymbol{m} = 0, \tag{S82b}$$

where, again, $\eta = \gamma_\perp/\gamma_\parallel$. (The component of the force balance perpendicular to the surface determines the reaction force $\boldsymbol{R}$.) Together with the condition of inextensibility, $\boldsymbol{r}_s\cdot\boldsymbol{r}_s = 1$, these equations determine $\theta(s,t)$, $\phi(s,t)$, $\Lambda(s,t)$ from the boundary [S9]

$$\boldsymbol{r}_{ss}(0,t)\cdot\boldsymbol{t}(0,t) = 0, \qquad \boldsymbol{r}_{ss}(0,t)\cdot\boldsymbol{m}(0,t) = 0, \tag{S83a}$$
$$\boldsymbol{r}_{ss}(\ell,t)\cdot\boldsymbol{t}(\ell,t) = 0, \qquad \boldsymbol{r}_{ss}(\ell,t)\cdot\boldsymbol{m}(\ell,t) = 0, \tag{S83b}$$

and

$$\bigl[\boldsymbol{r}_{sss}(0,t) + \Lambda(0,t)\boldsymbol{r}_s(0,t)\bigr]\cdot\boldsymbol{t}(0,t) = f, \tag{S83c}$$
$$\bigl[\boldsymbol{r}_{sss}(0,t) + \Lambda(0,t)\boldsymbol{r}_s(0,t)\bigr]\cdot\boldsymbol{m}(0,t) = 0, \tag{S83d}$$
$$\bigl[\boldsymbol{r}_{sss}(\ell,t) + \Lambda(\ell,t)\boldsymbol{r}_s(\ell,t)\bigr]\cdot\boldsymbol{t}(\ell,t) = a\boldsymbol{r}_t(\ell,t)\cdot\boldsymbol{t}(\ell,t), \tag{S83e}$$
$$\bigl[\boldsymbol{r}_{sss}(\ell,t) + \Lambda(\ell,t)\boldsymbol{r}_s(\ell,t)\bigr]\cdot\boldsymbol{m}(\ell,t) = \boldsymbol{r}_t(\ell,t)\cdot\boldsymbol{m}(\ell,t), \tag{S83f}$$

where, again, $a = \zeta_\parallel/\zeta_\perp$, and similarly to those that apply in the planar case.

#### B. Asymptotic expansion

From Eq. (S75a) with $q = p$, the position vector of a point on the germband midline is

$$\boldsymbol{r} = p\bigl(\cos\theta\cos\phi, \sin\theta, \cos\theta\sin\phi\bigr), \tag{S84a}$$

which implies that

$$\boldsymbol{n} = \bigl(\cos\theta\cos\phi, \sin\theta, \cos\theta\sin\phi\bigr), \tag{S84b}$$
$$\boldsymbol{t} = \boldsymbol{r}_s = p\bigl(-\theta_s\sin\theta\cos\phi - \phi_s\cos\theta\sin\phi, \theta_s\cos\theta, \phi_s\cos\theta\cos\phi - \theta_s\sin\theta\sin\phi\bigr), \tag{S84c}$$
$$\boldsymbol{m} = p\bigl(\theta_s\sin\phi - \phi_s\cos\theta\sin\theta\cos\phi, \phi_s\cos^2\theta, -\theta_s\cos\phi - \phi_s\cos\theta\sin\theta\sin\phi\bigr). \tag{S84d}$$

Inserting these expressions into the governing equations (S82) and boundary conditions (S83) yields explicit equations for

$\theta(s,t), \phi(s,t), \Lambda(s,t)$, but these are singularly unwieldy. However, to analyse the stability of the straight germband midline, it suffices to derive equations that describe small perturbations away from the straight germband midline. We therefore introduce a small parameter $\varepsilon \ll 1$ and an asymptotic expansion

$$\theta(s,t) = \theta_0(s,t) + \varepsilon\Theta(s,\tau) + O(\varepsilon^2), \tag{S85a}$$

$$\phi(s,t) = \varepsilon\Phi(s,t) + O(\varepsilon^2), \tag{S85b}$$

$$\Lambda(s,t) = \Lambda_0(s,t) + \varepsilon\ell(s,t) + O(\varepsilon^2), \tag{S85c}$$

in which $\theta_0(s,t)$ is given by Eq. (S79b). Correspondingly, we shall assume that $f = O(1)$. With this expansion, the condition of inextensibility becomes

$$1 = \boldsymbol{t}\cdot\boldsymbol{t} = 1 + 2p\varepsilon\Theta_s + O(\varepsilon^2) \quad\Rightarrow\quad \Theta(s,t) = \Theta(t). \tag{S86}$$

Then, from Eq. (S82a),

$$0 = -\left(\frac{u}{\eta} + \frac{\partial\Lambda_0}{\partial s}\right) + O(\varepsilon) \quad\Rightarrow\quad \Lambda_0(s,t) = -\frac{u}{\eta} + C(t), \tag{S87}$$

where $C(t)$ is a "constant" of integration and where $u$ is still undetermined. Finally, Eq. (S82b) gives, at order $O(\varepsilon)$, a partial differential equation for $\Phi(s,t)$ that simplifies to

$$\begin{aligned}\Phi_t = &\left\{u - \frac{2\left[p^2(su - f\eta) + \eta\right]}{p^3\eta}\tan\left(\frac{s+ut}{p}\right)\right\}\Phi_s \\ &+ \frac{p^2(su - f\eta) + 5\eta}{p^2\eta}\Phi_{ss} \\ &+ \frac{4}{p}\tan\left(\frac{s+ut}{p}\right)\Phi_{sss} \\ &- \Phi_{ssss}.\end{aligned} \tag{S88}$$

Now the first of Eqs. (S83a) and (S83b) are satisfied to order $O(\varepsilon^2)$, while the second of these equations yield two boundary conditions for $\Phi(s,t)$, viz.,

$$2\Phi_s(0,t)\sin\left(\frac{ut}{p}\right) - p\Phi_{ss}(0,t)\cos\left(\frac{ut}{p}\right) = 0, \tag{S89a}$$

and

$$2\Phi_s(\ell,t)\sin\left(\frac{\ell+ut}{p}\right) - p\Phi_{ss}(\ell,t)\cos\left(\frac{\ell+ut}{p}\right) = 0. \tag{S89b}$$

The remaining two boundary conditions required to solve Eq. (S88) are provided by Eqs. (S83d) and (S83f) which give, at order $O(\varepsilon)$,

$$\left[\frac{2}{p}\Phi_s(0,t) - p\Phi_{sss}(0,t)\right]\cos\left(\frac{ut}{p}\right) + 3\Phi_{ss}(0,t)\sin\left(\frac{ut}{p}\right) = 0, \tag{S89c}$$

and

$$\left[p\Phi_t(\ell,t) + \left(\frac{2}{p} - pu\right)\Phi_s(\ell,t) - p\Phi_{sss}(\ell,t)\right]\cos\left(\frac{\ell+ut}{p}\right) + 3\Phi_{ss}(\ell,t)\sin\left(\frac{\ell+ut}{p}\right) = 0. \tag{S89d}$$

Meanwhile, Eq. (S83c) yields, at order $O(1)$, $C(t) = f + p^{-2}$. Finally, Eq. (S83e) determines, at order $O(1)$,

$$u = \frac{\eta f}{\ell + a\eta}. \tag{S90}$$

However, Eq. (S88) is not separable due to its explicit dependence on time. In the "growing" model discussed above, this explicit dependence only appeared via the sum $\ell_0 + t$, and so the equation could be made separable by the slow-growth assumption $t \ll \ell_0$. By contrast, here, the explicit time dependence enters via the sum $s + ut$, with $s \in [0,\ell]$ and $t \geqslant 0$, so there is no similar assumption to be made. Hence Eq. (S88) does not allow a normal-mode analysis.

For this reason, stability analysis of the "pushed" model on the surface of a sphere is beyond the scope of this paper.

This makes a slightly more general point, that this kind of instability within a curved surface is rather difficult to analyse. Indeed, even Euler buckling within curved surfaces displays rich additional behaviour compared to the plane [S10].

---


[S1] G. De Canio, E. Lauga, and R. E. Goldstein, Spontaneous oscillations of elastic filaments induced by molecular motors, J. R. Soc. Interface **14**, 20170491 (2017).

[S2] The torque-free boundary conditions are stated in a form equivalent to Eqs. (S9) in Refs. [S1, S3], for example, but this is actually slightly subtle: From Eq. (S6), $\boldsymbol{r}_{ss}\cdot\delta\boldsymbol{r}_s = 0$ at a free boundary. The components of the variation $\delta\boldsymbol{r}_s$ are not independent however, because $\boldsymbol{r}_s\cdot\delta\boldsymbol{r}_s = 0$ from varying the constraint $\boldsymbol{r}_s\cdot\boldsymbol{r}_s = 1$. In the plane, this implies that $\boldsymbol{r}_s\times\boldsymbol{r}_{ss} = \boldsymbol{0}$, but not $\boldsymbol{r}_{ss} = \boldsymbol{0}$. This is of no consequence for the subsequent analysis, because, using $\boldsymbol{r}_s = \boldsymbol{t}$, $\boldsymbol{r}_{ss} = -\kappa\boldsymbol{n}$, $\boldsymbol{r}_s\times\boldsymbol{r}_{ss} = -\kappa\boldsymbol{t}\times\boldsymbol{n}$, so $\kappa = 0$ since $\boldsymbol{t}\perp\boldsymbol{n}$, which is equivalent to boundary conditions (S17).

[S3] Y. Man and E. Kanso, Morphological transitions of axially-driven microfilaments, Soft Matter **15**, 5163 (2019).

[S4] Y. Fily, P. Subramanian, T. M. Schneider, R. Chelakkot, and A. Gopinath, Buckling instabilities and spatio-temporal dynamics of active elastic filaments, J. R. Soc. Interface **17**,

20190794 (2020).

[S5] E. Lauga, *The Fluid Dynamics of Cell Motility*, Cambridge Texts in Applied Mathematics (Cambridge University Press, Cambridge, UK, 2020) Chap. 6, pp. 77–96.

[S6] D. Cholakova, M. Lisicki, S. K. Smoukov, S. Tcholakova, J. Chen, G. D. Canio, E. Lauga, and N. Denkov, Rechargeable self-assembled droplet microswimmers driven by surface phase transitions, Nat. Phys. **17**, 1050 (2021).

[S7] M. Abramowitz and I. A. Stegun, Handbook of mathematical functions (National Bureau of Standards, Washington, DC, 1964) Chap. 17, pp. 587–626.

[S8] M. P. Do Carmo, *Differential geometry of curves and surfaces* (Prentice-Hall, 1976) Chap. 4.4, pp. 238–264.

[S9] These boundary conditions suffer from a subtlety similar to that discussed in footnote [S2], but we will not discuss this in detail because we will not anyway solve these equations.

[S10] S. Zhao and P. A. Haas, Euler buckling on curved surfaces, Phys. Rev. Lett. **135**, 247201 (2025).
