## Supplementary Figures for "“Embryo-eggshell interaction counteracts chiral bias in early *Drosophila* morphogenesis”"

### Supplemental Figures

#### Suppl. Fig. S1

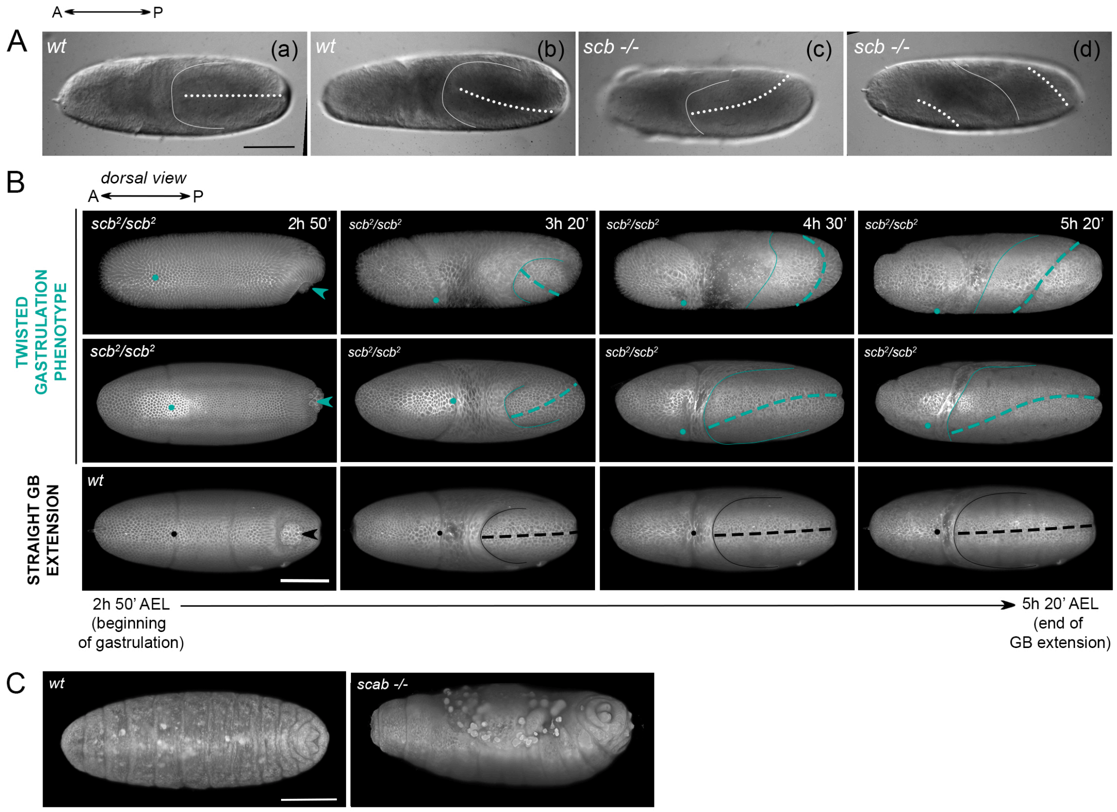


**Suppl. Fig. S1: The curved germ band extension phenotype in wild-type and *scab* mutants**

A) Brightfield recording of germ band extension in wild-type (wt, *Oregon R*) and *scab* loss of function (*scab* $^{-/-}$, *scb*$^2$/*scb*$^2$) embryos. (a) Straight GB extension, (b), (c) curved GB extension, (d) twisted gastrulation phenotype. Scale bar: 100 µm.

B) 3D rendering of a light sheet microscopy recording of germ band extension in wild-type (wt, homozygous for the membrane marker RGAP43::mCherry) and *scab* loss of function (*scb*$^2$/*scb*$^2$) embryos. Scale bar: 100 µm.

C) 3D rendering of embryos expressing RGAP43::mCherry fluorescent membrane marker at dorsal closure, showing a wild-type and a *scab* loss of function phenotype (*scb*$^2$/*scb*$^2$). Scale bar: 100 µm.

### Suppl. Fig. S2

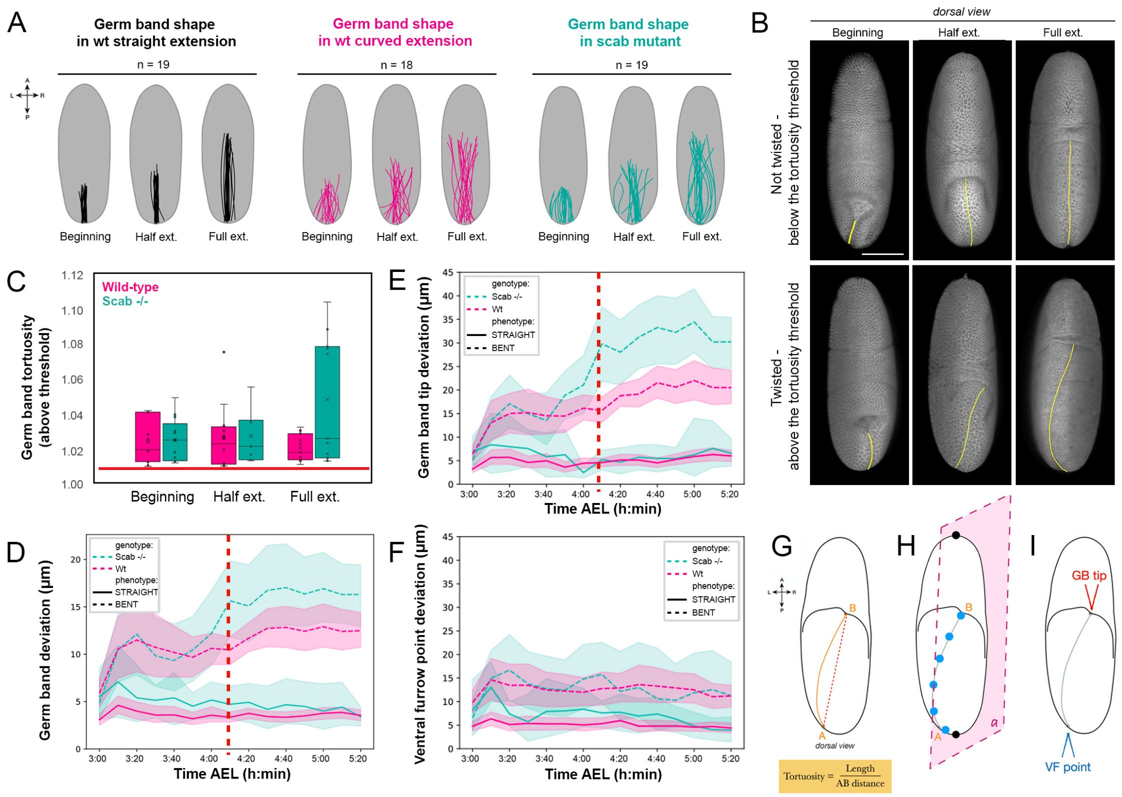


### Suppl. Fig. S2: 2D and 3D analysis of germ band deformation

A) Diagrams representing the variability of the shape of the germ midline at selected timepoints observed in subsets of developing embryos displaying the canonical (black) or altered (magenta, green) germ band extension phenotypes. Dorsal views.

B) 3D renderings, at the timepoints of interest (Fig. 1A), of representative embryos imaged by light sheet microscopy that are above and below the tortuosity threshold for curved extension. The yellow line highlights the shape of the germ band midline. Dorsal views. Scale bar: 100 µm.

C) Boxplot of the distribution of tortuosity values above the threshold of T = 1.01 (red line) defining the altered extension phenotype, at the three timepoints of interest in the wild-type (*RGAP43::mCherry*) and *scab* loss of function ($scb^2/scb^2$) samples (the samples are significantly different in all three time points, see Fig. 1F).

D) Plot of the 3D deviation from straightness values over time in *RGAP43::mCherry* and $scb^2/scb^2$ mutant embryos. A threshold of 6.9 has been applied to separate each data group in two classes: "straight" (full line) and "bent" (dashed line) for visualization purposes. The vertical red dashed line indicates the time point after which the difference between the two sample groups becomes statistically significant (see Fig. 1D).

E) Plot of the 3D deviation from straightness values at the tip of the germ band over time in *RGAP43::mCherry* and $scb^2/scb^2$ mutant embryos. A threshold of 6.9 has been applied to separate each data group in two classes: "straight" (full line) and "bent" (dashed line) for visualization purposes. The vertical red dashed line indicates the time point after which the difference between the two sample groups becomes statistically significant (see Fig. 1G).

F) Plot of the 3D deviation from straightness values at the ventral furrow point over time in *RGAP43::mCherry* and $scb^2/scb^2$ mutant embryos. A threshold of 6.9 has been applied to separate each data group in two classes: "straight" (full line) and "bent" (dashed line) for visualization purposes.

G) Schematic illustrating the calculation of the tortuosity of the germ band midline.

H) Schematic illustrating the definition of the deviation from straightness of the germ band midline: the deviation from straightness minimizes the distance of specific points along the germ band midline (blue dots) to the planes

*a* (pink) passing through the center of the embryo along its longitudinal axis, identified by the two black dots (Supplemental Methods).
I) Schematic definition of the two points of interest analyzed separately: “GB tip” indicates the tip of the germ band and “VF point” indicates the point where the germ band midline emerges from the ventral side of the embryo.

### Suppl. Fig. S3

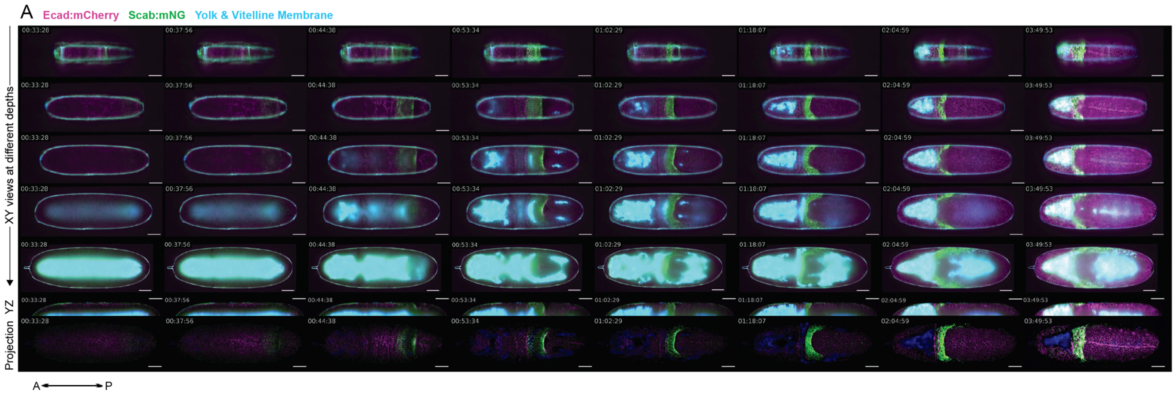


**Suppl. Fig. S3: Scab::mNG expression pattern during germ band extension**

Ecad::mCherry (magenta), Scab::mNG (green) and autofluorescence of yolk and vitelline membrane (cyan) signals. Sagittal YZ views at different depths from the raw data and local Z-projections of the processed data. Scale bars: 50 µm.

**Suppl. Fig. S4**

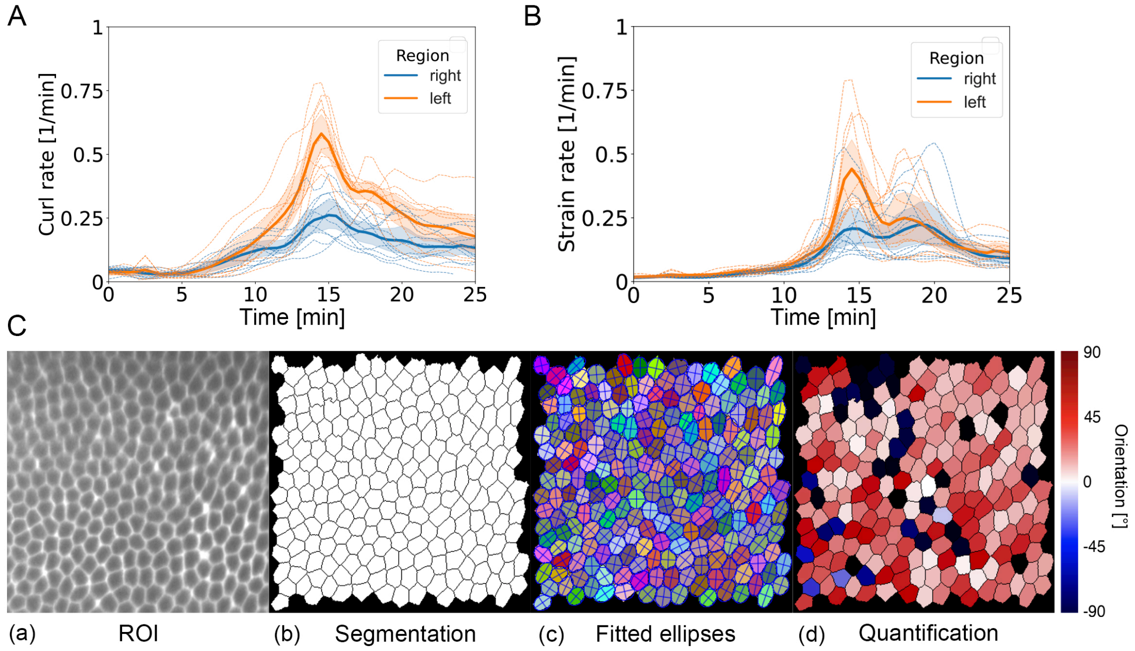


**Suppl. Fig. S4: Embryo asymmetry on the tissue and cellular level**

A) Curl rates in the left (orange) or right (blue) ROIs in Fig. 4G against time $t$, where $t = 0$ is the end of cellularization. Dotted lines: data for each of n = 10 embryos, solid lines: averages

B) Strain rates in the left (orange) or right (blue) ROIs in Fig. 4H against time $t$, where $t = 0$ is the end of cellularization. Dotted lines: data for each of n = 10 embryos, solid lines: averages

C) Analysis of cellular geometric features: (a) analyzed Region Of Interest (ROI); (b) watershed segmentation with removal of objects at the edges; (c) labelled objects fitted by ellipses; (d) color-coded objects according to the orientation with respect to the L-R axis.

### Suppl. Fig. S5

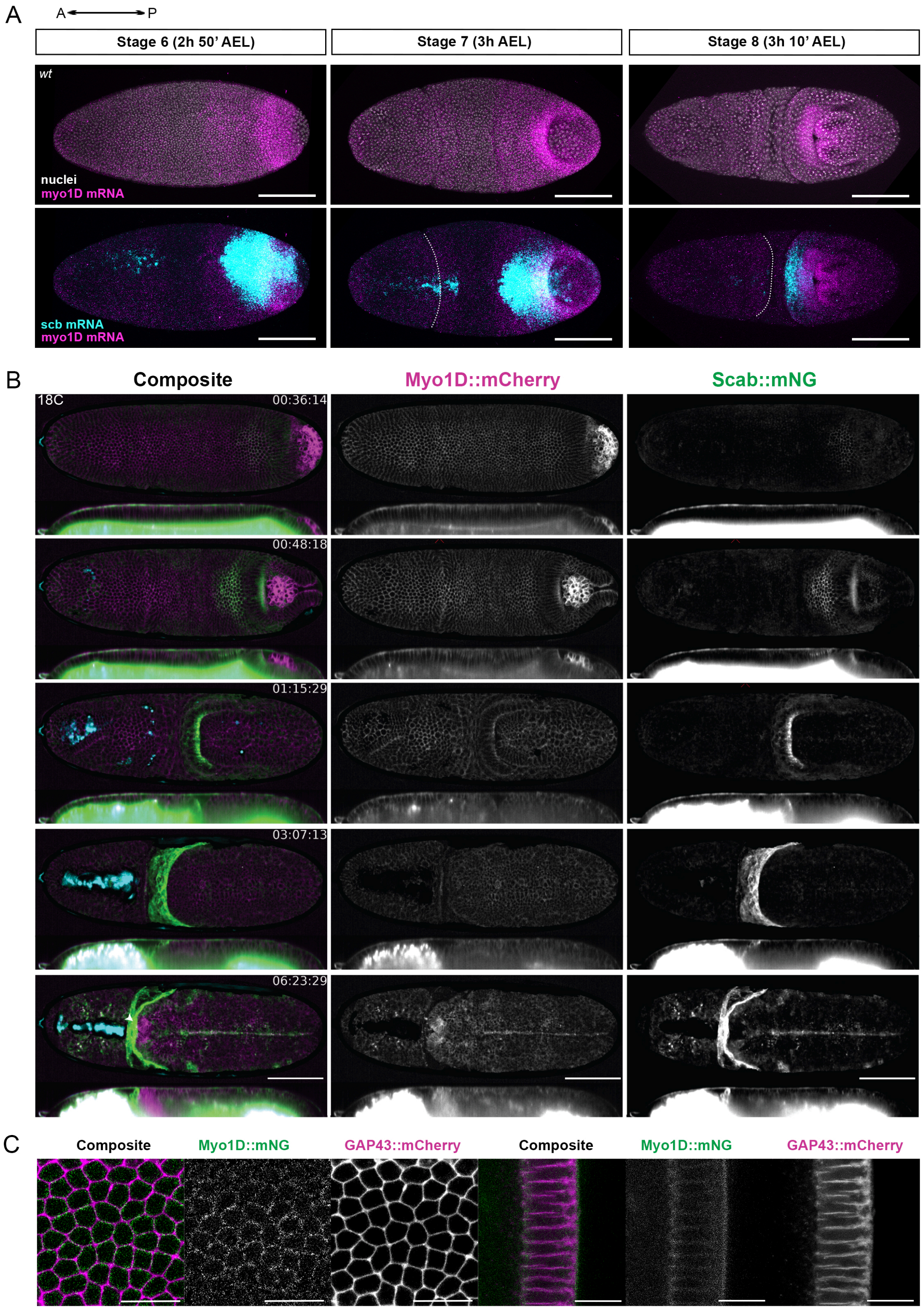

A
A P
Stage 6 (2h 50' AEL)
Stage 7 (3h AEL)
Stage 8 (3h 10' AEL)
wt
nuclei
myo1D mRNA
scb mRNA
myo1D mRNA
B
Composite
Myo1D::mCherry
Scab::mNG
18C
00:36:14
00:48:18
01:15:29
03:07:13
06:23:29
C
Composite
Myo1D::mNG
GAP43::mCherry
Composite
Myo1D::mNG
GAP43::mCherry

**Suppl. Fig. S5: *Myo1D* and *scab* RNA and protein expression in embryogenesis**

A) Expression of *myo1D* (magenta) and *scab* (cyan) mRNA in fixed wild-type embryos at selected timepoints. Stage 6: before gastrulation, stage 7: during gastrulation, stage 8: during the fast phase of germband extension in Fig. 1 (Campos-Ortega and Hartenstein, 1985). Nuclei are shown in gray. Top row: lateral view; bottom row: dorsal view; white dotted lines: position of the cephalic furrow.

B) Live light sheet microscopy recording of the expression pattern of transgenic Myo1D::mCherry (magenta) and Scab::mNG (green) reporters at selected timepoints representative of different phases of germ band extension. Each timepoint is shown both from the dorsal view and in a cross section along the longitudinal axis. Scale bars: 100 µm.

C) High-resolution live confocal recording of Myo1D::mNG (green) reporter line at cellularization, showing colocalization of Myo1D protein with the cellularization front. The cell membrane is labelled by RGAP43::mCherry fluorescent marker (magenta). Dorsal (left) and lateral (right) views. Scale bars: 100 µm.

**Suppl. Fig. S6**

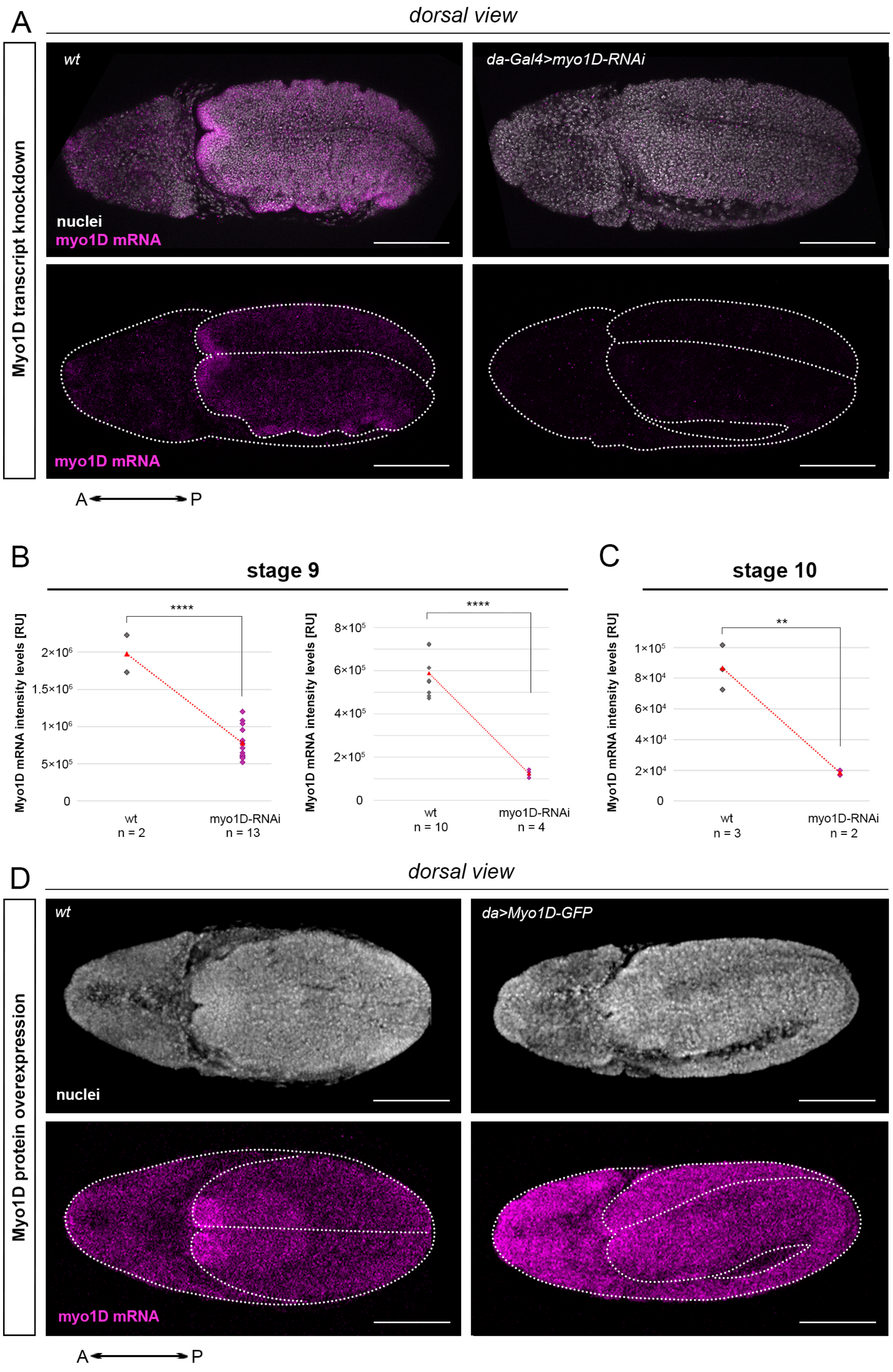

A
dorsal view
Myo1D transcript knockdown
wt
da-Gal4>myo1D-RNAi
nuclei
myo1D mRNA
myo1D mRNA
A P
B
stage 9
Myo1D mRNA intensity levels [RU]
****
2×10^6
1.5×10^6
1×10^6
5×10^5
0
wt
n = 2
myo1D-RNAi
n = 13
Myo1D mRNA intensity levels [RU]
****
8×10^5
6×10^5
4×10^5
2×10^5
0
wt
n = 10
myo1D-RNAi
n = 4
C
stage 10
Myo1D mRNA intensity levels [RU]
**
1×10^5
8×10^4
6×10^4
4×10^4
2×10^4
0
wt
n = 3
myo1D-RNAi
n = 2
D
dorsal view
Myo1D protein overexpression
wt
da>Myo1D-GFP
nuclei
myo1D mRNA
A P

**Suppl. Fig. S6: Quantification of *myo1D* transcript in knockdown and overexpression experiments**

A) Expression of *myo1D* transcript in wild-type (*Oregon R*) and *da-Gal4; UAS-myo1D-RNAi* embryos at the end of germ band extension. Myo1D transcript is indicated in magenta, nuclei are in gray. Lateral views. Scale bars: 100 µm.

B) Reduction of *myo1D* mRNA signal intensity in conditions of *myo1D* RNAi knockdown at stage 9 (both replicates, $p < 0.0001$. Two-sample one-tailed T-test assuming unequal variances, p-values in Suppl. Table XI.

C) Reduction of *myo1D* mRNA signal intensity in conditions of *myo1D* RNAi knockdown at stage 10: $p < 0.001$. Two-sample one-tailed T-test assuming unequal variances, p-values in Suppl Table XI.

D) Expression of *myo1D* transcript in wild-type (*Oregon R*) and *da-Gal4; UAS-Myo1D::GFP* embryos at the end of germ band extension. Myo1D transcript is indicated in magenta, nuclei are in gray. Lateral views. Scale bars: 100 µm.
